# The OCT2 Transporter Regulates Dopamine D1 Receptor Signaling at the Golgi Apparatus

**DOI:** 10.1101/2020.11.23.394445

**Authors:** Natasha M. Puri, Giovanna R. Romano, Ting-Yu Lin, Quynh N. Mai, Roshanak Irannejad

**Author notes:** These authors contributed equally (Natasha M. Puri, Giovanna R. Romano).

## Abstract

Dopamine is the main catecholamine in the brain and the kidney, where it controls a number of physiological functions such as locomotion, cognition, emotion, endocrine regulation and renal function. As a membrane impermeant hormone and neurotransmitter, dopamine is thought to signal by binding and activating dopamine receptors, members of the G protein couple receptor (GPCR) family, only on the plasma membrane. Here, using novel nanobody-based biosensors, we demonstrate for the first time that the dopamine D1 receptor (D1DR), the primary mediator of dopaminergic signaling in the brain and kidney, not only functions on the plasma membrane but becomes activated at the Golgi apparatus in the presence of its ligand. We present evidence that activation of the Golgi pool of D1DR is dependent on Organic Cation Transporter 2 (OCT2), a dopamine transporter, providing an explanation for how the membrane impermeant dopamine accesses subcellular pools of D1DR. We further demonstrate that dopamine activates Golgi-D1DR in the striatal medium spiny neurons (MSN) and this activity depends on OCT2 function. We also introduce a new approach to selectively interrogate compartmentalized D1DR signaling by inhibiting Gαs coupling, using a nanobody-based chemical recruitment system. Using this strategy, we show that Golgi-localized D1DRs regulate cAMP production and mediate local protein kinase A activation. Together, our data suggest that spatially compartmentalized signaling hubs are previously unappreciated regulatory aspects of D1DR signaling. Our data also provide further evidence for the role of transporters in regulating subcellular GPCR activity.

## Introduction

Dopamine (DA) is a major hormone and neurotransmitter that regulates a wide range of physiological responses including reward-motivated behavior, aversion, cognition, and motor control in the central nervous system, CNS(1-3). DA also regulates physiological responses in non-CNS tissues such as sodium secretion in the kidney (4). All cellular actions of DA are mediated by Dopamine receptors, a class of GPCR. Several pathological conditions such as Parkinson’s disease, schizophrenia and addiction are due to dysregulation of the neuronal dopaminergic signaling pathway, while hypertension has been attributed to impaired renal dopaminergic signaling (5, 6). Dopamine receptor antagonists have been developed with the goal of blocking hallucinations and delusions that occur in schizophrenic patients, whereas DA receptor agonists are used to alleviate the moving defects of Parkinson’s disease (5-7).

In both the CNS and the kidney, DA is produced locally. There are five subtypes of Dopamine receptors, D1, D2, D3, D4 and D5, that are classified as D1-class receptors (D1 and D5) or D2-class receptors (D2, D3 and D4) (8, 9). The D1-class (D1 and D5 receptors) are primarily coupled to Gαs/olf proteins and stimulate the activity of adenylyl cyclase, AC, leading to the production of the second messenger cyclic AMP, cAMP (10). In contrast, the D2 class (D2S, D2L, D3 and D4 receptors) are associated with Gαi/o proteins and inhibit cAMP production (10). D1 Dopamine Receptors (D1DRs) are highly expressed in the CNS where they underlie major brain functions such as locomotion, learning and memory, attention, impulse control, and sleep (4). Additionally, D1DRs in the kidney regulate trafficking of sodium ATPase and transporters, thereby affecting renal function (11, 12).

Impermeable agonists such as DA have been thought for a long time to activate D1DR only at the plasma membrane. Like many GPCRs, receptor removal from the cell surface by endocytosis has been described as a mechanism that attenuates cellular signaling (13-15). As such, efforts at modulating DA signaling as a therapeutic strategy for various pathophysiological conditions have only taken into consideration the consequences of signaling by plasma membrane localized DA receptors(16-18). However, evidence from the past decade suggests that for some GPCRs endocytosis might in fact activate a second phase of acute or prolonged Gαs-mediated cAMP response from the endosomes (19-29). Recent studies further support this notion by providing evidence that cAMP generation by activated receptors at the endosome are necessary in regulating transcriptional responses that are distinct from those elicited by activation of the plasma membrane receptor pool (30-35).

Most of the receptors that have been shown to exhibit a second phase of signaling from internal compartments are primarily coupled to Gαs protein. cAMP diffusion is within the nanometer scale around phosphodiesterases at physiological conditions (36-38). Given this narrow range of diffusion, it has been difficult to explain how receptor activation solely on the plasma membrane results in the activation of downstream effectors at distant subcellular locations such as the endoplasmic reticulum, Golgi and the nucleus (37-39). As one explanation for this observation, we recently showed that activation of the Golgi-localized beta1 adrenergic receptors (β1AR) cause local production of cAMP by Golgi localized Gαs protein. Importantly, we demonstrated that a catecholamine transporter facilitates the transport of epinephrine, a membrane impermeant endogenous β1AR agonist, to the lumen of the Golgi and activates the Golgi pool of β1AR (40). The importance of generation of a local pool of cAMP by Golgi-localized β1AR was further supported by the fact that activated Golgi-β1ARs only, but not the plasma membrane pool, lead to PLCε activation at the perinuclear/Golgi membrane, which mediates hypertrophic responses in cardiomyocytes (40, 41).

Whether the need for local cAMP generation is unique to cell types or specific GPCRs is not well understood. The lack of cAMP mobility in cells becomes prominent in larger cells with higher membrane compartmentation, which will further create physical barriers for cAMP diffusion. Considering the high degree of membrane compartmentation of neurons and proximal tubules of the kidney, the two main cells types that express D1DRs, we wondered whether D1DR signaling is also compartmentalized. Here, using a conformational sensitive nanobody that recognizes activated D1DR, we show that the pre-existing pool of D1DR that is localized to the Golgi membrane is activated upon stimulation with extracellular DA. In addition to several cell lines, here we demonstrate that Golgi-localized D1DR signaling is also a feature of primary striatal medium spiny neurons (MSNs). The D1DR-expressing MSNs of striatum are well established to play significant roles in motivation, aversion and reward (3, 42-45). Furthermore, we demonstrate that OCT2 facilitates the transport of DA to the Golgi-localized D1DR and regulates its local activity at the Golgi. We further show that OCT2 has a distinct expression pattern in the kidney and specific regions of the brain including the MSNs, where D1DRs are endogenously expressed. Thus, our findings reveal that DA can activate D1DR signaling at the Golgi and point to a novel role for OCT2 as a factor that determines which cell types exhibit DA-mediated subcellular signaling.

## Results

### Nanobody-based conformational sensitive biosensors detect active D1DR and Gs protein at subcellular membranes

We have previously shown that a single-domain camelid antibody, nanobody 80 (Nb80), originally developed to stabilize an active conformation of beta 2 adrenergic receptor (β2AR) for crystallography purposes (46), can be repurposed as a conformational biosensor to detect activated β2AR and β1AR in living cells (23, 40). Through directed evolution on Nb80, a high-affinity nanobody (Nb6B9) was generated that stabilizes the active conformation of epinephrine bound β2AR (47). Given that β2AR/Nb6B9 binding sites are highly conserved among other aminergic receptors such as β1AR and D1DR (Supplementary Figure 1)(46), we reasoned that this nanobody could also be used as a conformational sensitive biosensor to detect activated D1DR in real time and in living cells (Figure 1a). In HeLa cells expressing Snap-tagged D1DR, Nb6B9 fused to GFP (Nb6B9-GFP) was diffuse throughout the cytoplasm (Figure 1b, 0min). Upon stimulation of D1DR expressing HeLa cells with 10μM DA, Nb6B9-GFP was rapidly recruited first to the plasma membrane and shortly after to the Golgi apparatus (Figure 1b, 2min, Supplementary Movie 1). This data suggests that the D1DR Golgi pool is activated in response to extracellular DA addition. Interestingly, stimulation of 10μM DA in HEK293 cells expressing D1DR resulted in the recruitment of NB6B9-GFP only at the plasma membrane (Figure 1b, lower panel, 2min, Supplementary Movie 2). However, in both cell types, SKF81297, a selective membrane permeant D1DR agonist, activated both receptor pools (Supplementary Figure 2 and Supplementary movie 3 and 4). In addition to the Golgi recruitment and consistent with a previous report (22), Nb6B9 was also found to colocalize with D1DR at the endosomes, at a later time after agonist addition, indicating an active pool of D1DR at endosomes (Supplementary Movie 2). We further used mini-Gs protein, a more general biosensor for Gs-coupled GPCRs (48), to show that the active pool of D1DR at the plasma membrane, endosomes, and the Golgi can also be detected by mini-Gs recruitment to these locations (Supplementary Figure 3).

**Figure 1.**
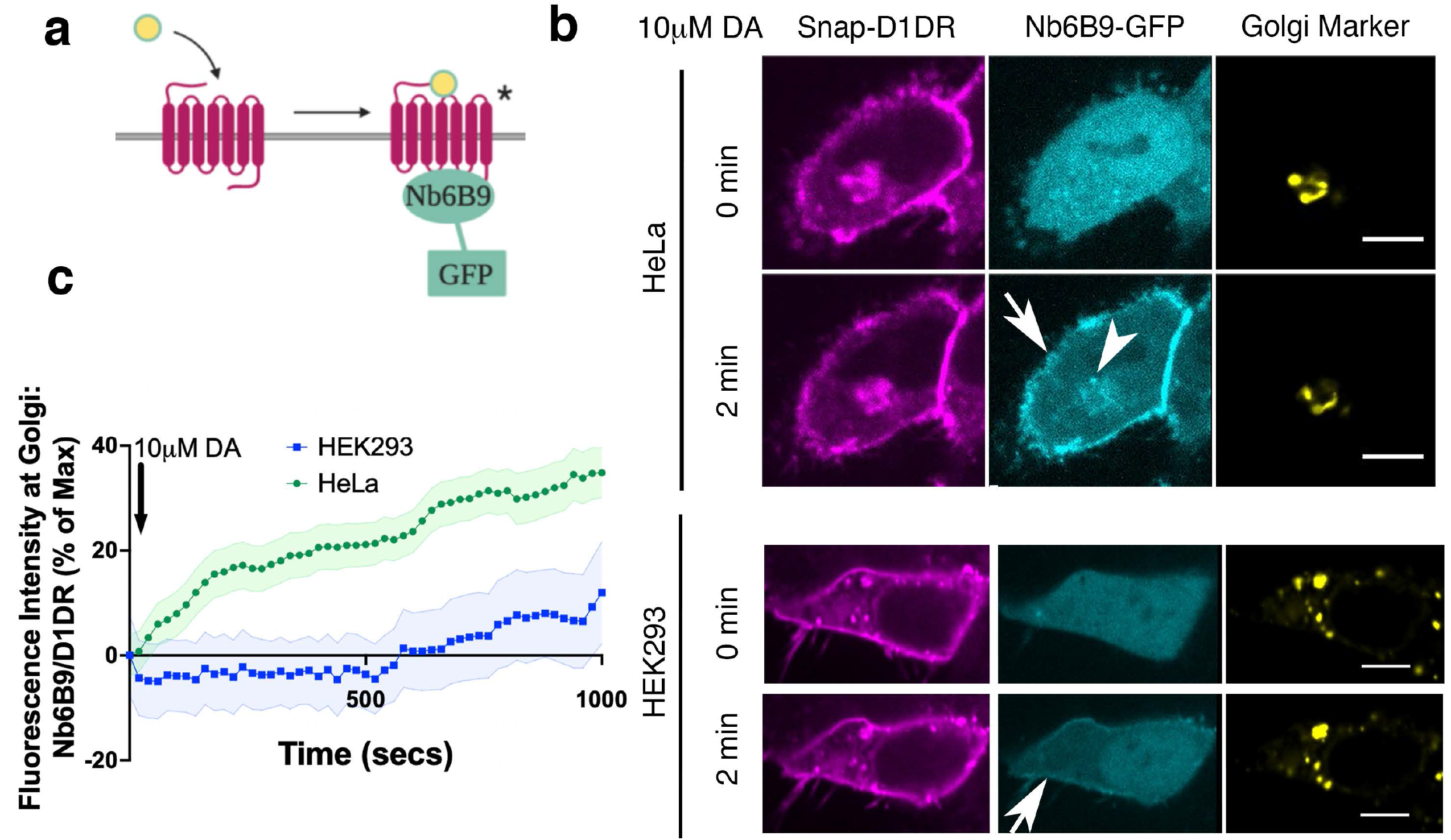
Conformational biosensor detects activated D1DR at the plasma membrane and the Golgi upon Dopamine stimulation. **a**. Nb6B9 binds to the receptor exclusively in its active conformation. We fused Nb6B9 to GFP and used it as conformational biosensor for D1DR. **b**. Confocal images of representative D1DR-expressing HeLa (top panel) and HEK293 cells (lower panel) with Nb6B9-GFP and GalT-mRFP expression before and after 10μM dopamine addition. Stimulation with 10μM dopamine results in recruitment of Nb6B9 to active D1DR at plasma membrane and Golgi in HeLa (*n* = 15 cells, Pearson’s coefficient = 0.63, respectively, 5 biological replicates); 10μM DA treatment only activates plasma membrane localized D1DR in HEK 293 cells (*n* = 17 cells, Pearson’s coefficient = 0.15, 5 biological replicates). Right panels show merged images of Nb6BP and the Golgi marker. Arrows indicate active D1DR at plasma membrane; Arrowhead indicates active D1DR at Golgi membrane; Scale bar = 10μm **c**. Quantification of D1DR activity at Golgi in HeLa and HEK293 cells; Normalized fluorescence intensity of Nb6B9 at Golgi relative to D1DR at Golgi labeled with Snap-tagged-D1DR.

To investigate whether activated D1DR can couple to G protein and elicit a G-protein mediated response at the Golgi, we took advantage of another nanobody-based biosensor, Nb37-GFP. We previously used Nb37-GFP to detect transiently active β1AR/Gs and β2AR/Gs complexes at the Golgi and endosomes, respectively (23, 40). Similar to what we previously observed with β1AR, Nb37 was recruited to the plasma membrane and the Golgi upon stimulation with DA, suggesting that the D1DR Golgi pool is able to couple to G protein and activate it (Supplementary Figure 4b and c). Together, these findings suggest a distinct spatiotemporal regulation of D1DR signaling at the plasma membrane and the Golgi membranes.

### OCT2 facilitates the transport of Dopamine to the Golgi-localized D1DR

These observations raised the key question of how DA, a hydrophilic/membrane impermeant monoamine, can access the Golgi pool of D1DR. The first clue came from the observation that DA can activate Golgi-D1DR in HeLa cells but not HEK293 cells (Figure 1b, Supplementary Movie 1 and 2) whereas SKF81297, a hydrophobic/membrane permeant agonist, can activate the Golgi pool of D1DR in both cell types (Supplementary Figure 2, Supplementary Movie 3 and 4). These distinct effects of DA and SKF81297 are not based on their differential potency for activating D1DR, as they have comparative EC50 values in inducing cAMP responses (Supplementary Figure 5a). Moreover, D1DR activation at the Golgi is not dependent on receptor internalization, as inhibiting endocytosis by blocking dynamin did not block D1DR activation at the Golgi (Supplementary Figure 4a and c). We previously found that a membrane transporter, Organic Cation Transporters 3 (OCT3), facilitates epinephrine transport resulting in activation of the Golgi-localized β1AR. OCT3 is a member of the solute carrier (SLC) family 22, uptake 2 transporters that are electrogenic and transport catecholamines in a bidirectional manner. Importantly, OCT transporters are localized on the plasma membrane and intracellular compartments including nuclear envelope, thus they can transport catecholamines across the plasma membrane and across internal membrane compartments (40, 49). Therefore, we hypothesized that another OCT family transporter can similarly function in DA transport to allow for its delivery to the Golgi and for the activation of Golgi-localized D1DR pools.

There are three main OCTs that have largely overlapping distribution but distinct substrates(50-53). OCT3 facilitates the transport of epinephrine and norepinephrine (50). DA has been identified as a key endogenous substrate for another member of the SLC22A family, OCT2 (SLC22A2)(52-57). We thus asked whether OCT2 has a role in transporting DA to the Golgi membranes. We found significant OCT2 protein expression in HeLa cells as measured by western blotting, whereas no significant expression was detected in HEK293 cells (Supplementary Figure 5b). To test the role of OCT2 in DA transport, we first overexpressed OCT2-mApple in HEK293 cells and used Nb6B9-GFP to assess D1DR activation in live cells. By overexpressing OCT2-mApple in HEK293 cells, we found that Nb6B9-GFP could now be recruited to activated D1DR at the Golgi membranes. (Figure 2a and b, Supplementary Movie 5). Second, we inhibited OCT2 in HeLa cells, using a selective inhibitor (100 μM imipramine), and found that DA-mediated D1DR activation at the Golgi is inhibited. Moreover, SKF81297, a membrane permeant D1DR agonist that can diffuse across the membrane and does not require facilitated transport, can still access and activate Golgi-D1DR in imipramine-treated cells (Figure 2 c and d, lower panel and Supplementary Movie 6). Importantly, OCT2 but not OCT3 is required for DA transport and activation of Golgi-D1DR, as OCT3 inhibition did not have any effect on DA-mediated activation of D1DR Golgi pool (Figure 2d, top panel and Supplementary Figure 6). Altogether, these results suggest that OCT2 facilitates the transport of DA to the Golgi lumen where it then activates D1DR at the Golgi membranes.

**Figure 2.**
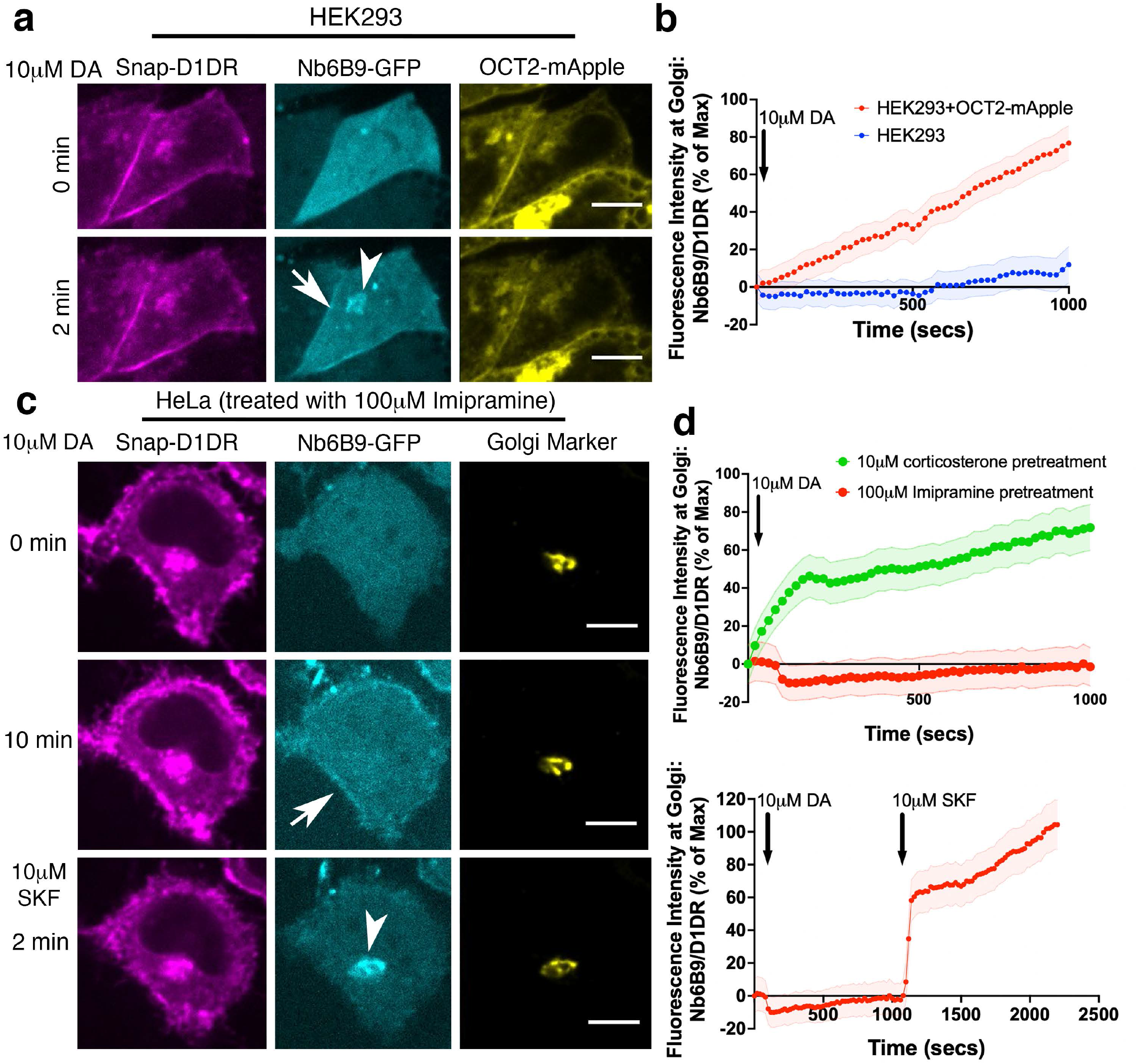
OCT2 facilitated DA transport to the Golgi-Localized D1DR. **a**. Representative HEK293 cell expressing Snap-D1DR, Nb6B9-GFP, and OCT2-mApple at indicated times after 10μM dopamine addition. Overexpression of OCT2 in HEK293 cells rescues Golgi-localized D1DR activation (*n* = 16 cells, Pearson’s coefficient = 0.67, 6 biological replicates). Arrow indicates active D1DR at plasma membrane; Arrowhead indicates active D1DR at Golgi membrane; Scale bar = 10μm. **b**. Quantification of Nb6B9-GFP recruitment at Golgi in HEK293 cells with or without OCT2 overexpression; Normalized fluorescence intensity of Nb6B9-GFP relative to Snap D1DR at Golgi. **c**. Representative HeLa cell expressing Snap-D1DR, Nb6B9-GFP, and GalT-mRFP expression pretreated with 100μM imipramine for 15 min, before and after 10μM dopamine addition. Inhibition of OCT2 blocks Golgi-localized D1DR activation but SKF81297 can still reach the Golgi membranes and activate D1DR Golgi pool (*n* = 16 cells, Pearson’s coefficient = 0.67, 6 biological replicates). Arrow indicates active D1DR at plasma membrane; Arrowhead indicates active D1DR at Golgi membrane; Scale bar = 10μm. **d**. Quantification of Nb6B9-GFP recruitment at Golgi upon 10μM DA stimulation in HeLa cells pretreated with OCT2 (100μM imipramine) or OCT3 (10μM corticosterone) inhibitors and after 10μM SKF81297 addition; Normalized fluorescence intensity of Nb6B9-GFP relative to Snap D1DR at Golgi (*n*= 30, 5 biological replicates).

### Regulation of dopamine-mediated activation of the Golgi-localized D1DR in striatal MSNs by OCT2

To investigate the role of OCT2 in D1DR signaling in physiologically relevant cell types, we measured OCT2 expression patterns in two main organs where D1DR are known to express and have function, the kidney and the brain. We found that OCT2 is highly expressed in the kidney and has a distinct expression pattern in different regions of the brain. OCT2 is highly expressed in the striatum and cortex, where D1DR is known to express and have function(58-60). However, OCT2 expression was negligible in the hippocampus and substantia nigra, another region where D1DR has known functions (Supplementary Figure 5c) (43, 54, 61, 62).

To determine the role of OCT2 in regulating a distinct pool of D1DR signaling in neurons, we isolated primary murine striatal MSNs, where OCT2 is expressed at high levels (Supplementary Figure 5c). Within the striatum, D1DR-expressing MSNs have been shown to play roles in DA-regulated processes such as motivation, aversion and reward seeking (3, 42-45). We detected endogenous D1DR on both the plasma membrane and the Golgi membranes in MSNs (Supplementary Figure 5d, top panel). Stimulating D1DR expressing MSNs with DA resulted in the recruitment of Nb6B9-GFP to both the plasma membrane and the Golgi membrane (Figure 3a, Supplementary Figure 5d lower panel, Supplementary Movie 7). Importantly, OCT2 inhibition resulted in the inhibition of DA-mediated Golgi-D1DR activation. By contrast, SKF81297, a membrane permeant D1DR agonist that does not require a transporter to reach the Golgi-D1DR pool, activated D1DR at the Golgi (Figure 3b and Supplementary Movie 8). These data demonstrate that Golgi-localized signaling by D1DRs occurs in a physiologically relevant cell type and that this signaling requires OCT2. Moreover, as there are cell types that express D1DR but not OCT2, our findings suggest that OCT2 serves as a factor that determines which cell types will exhibit both plasma membrane and Golgi-localized D1DR signaling under physiological conditions.

**Figure 3.**
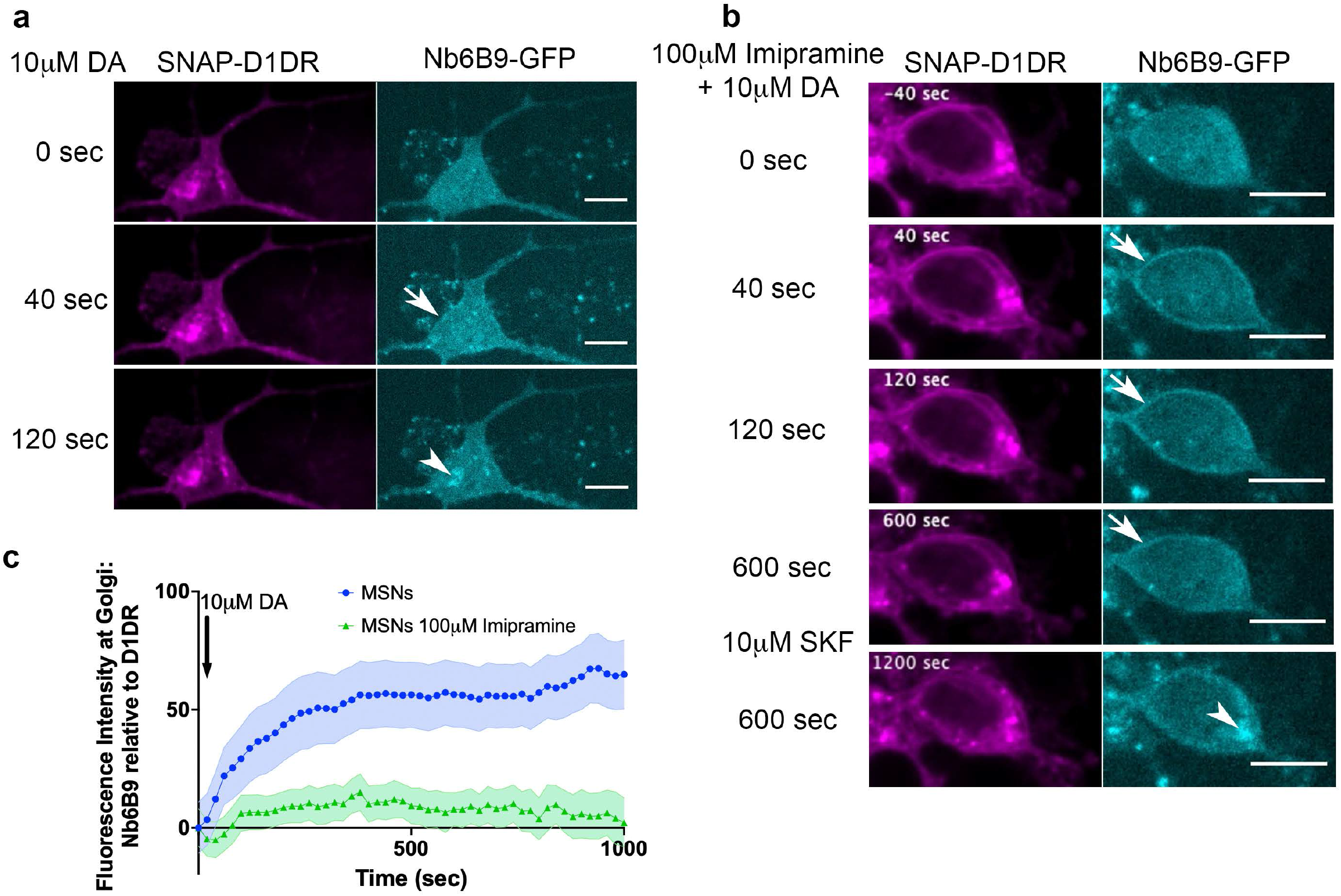
Regulation of dopamine-mediated activation of the Golgi-localized D1DR in striatal neurons by OCT2. **a**. Representative MSN expressing Snap-D1DR and Nb6B9-GFP at indicated times after 10μM dopamine addition. DA stimulates D1DR activation at the Golgi in MSNs (*n* = 22 cells, Pearson’s coefficient = 0.67, 6 biological replicates). Arrow indicates active D1DR at plasma membrane; Arrowhead indicates active D1DR at Golgi membrane; Scale bar = 10μm. **b**. Representative MSN cell expressing Snap-D1DR and Nb6B9-GFP, pretreated with 100μM imipramine for 15 min, before and after 10μM dopamine addition. Inhibition of OCT2 blocks Golgi-localized D1DR activation at MSN (*n* = 18 cells, Pearson’s coefficient = 0.38, 6 biological replicates) but SKF81297 can still reach the Golgi membranes and activate D1DR Golgi pool (*n* = 6 cells, Pearson’s coefficient = 0.75 4 biological replicates). Arrow indicates active D1DR at plasma membrane; Arrowhead indicates active D1DR at Golgi membrane; Scale bar = 10μm. **c**. Quantification of Nb6B9-GFP recruitment at Golgi upon 10μM DA stimulation in MSNs cells pretreated with OCT2 inhibitor; Normalized fluorescence intensity of Nb6B9-GFP relative to Snap D1DR at Golgi (*n*= 12 and 7, respectively, 5 biological replicates).

### Golgi and plasma membrane-localized D1DR both contribute to the cAMP response

Our data suggested that the plasma membrane and Golgi pools of D1DR both couple to the Gs protein. In addition to its presence at the plasma membrane, adenylyl cyclase has been reported to localize at the Golgi/perinuclear membranes (63, 64). We therefore asked whether D1DR/Gs complexes at both the plasma membrane and the Golgi activate Gs-mediated cAMP responses. To address this question, we utilized a rapamycin dimerization system composed of FK506-binding protein (FKBP) and FKBP-rapamycin binding domain of FRAP (FRB), to rapidly induce recruitment of Nb6B9 to specific membrane compartments. This makes it possible to specifically block D1DR/Gs coupling at each distinctly localized pool. We have previously shown that βARs nanobody, Nb80, which binds to the same region as G protein (46, 65), blocks either the plasma membrane or the Golgi-β1AR mediated cAMP responses when it is recruited locally to these compartments at high concentrations(40). This inhibition is likely due to steric occlusion of the Gαs protein. Using HEK293 cells expressing either FKBP at the plasma membrane or the Golgi with FRB fused to Nb6B9 (FRB-Nb6B9), we demonstrated that treatment with rapalog, a rapamycin analog, specifically targets Nb6B9 to either membranes (Figure 4 a-c). Upon stimulation with membrane permeant agonist SKF81297, Nb6B9 targeted to the plasma membrane disrupts plasma membrane-D1DR /G proteins coupling, while Golgi-D1DR is still able to elicit a cAMP response (Figure 4d). In turn, treatment with rapalog in cells expressing Golgi-targeted FKBP and FRB-Nb6B9 and subsequent stimulation with SKF81297 resulted in only activation of the plasma membrane-D1DR pool (Figure 4e). Importantly, rapalog treatment alone had no effect on the overall cAMP production elicited by Forskolin, a direct activator of adenylyl cyclase (Figure 4f). These data indicate that Golgi-localized D1DR is able to promote cAMP response.

**Figure 4.**
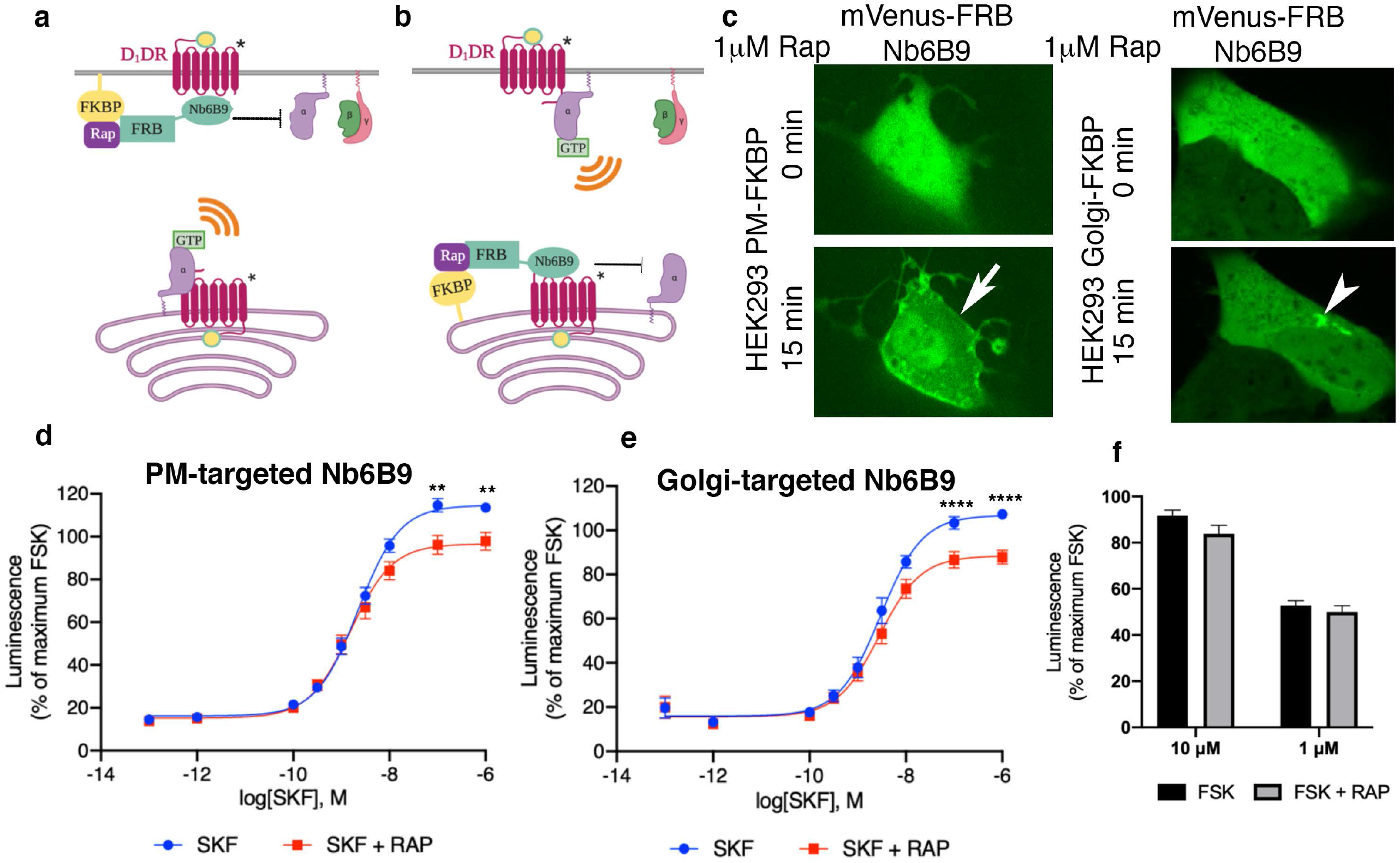
Both plasma membrane and Golgi-localized D1DR promote cAMP production. Model of blocking D1DR-Gs coupling at the plasma membrane (**a**) and Golgi membrane (**b**) after recruitment of mVenus-FRB-Nb6B9. FKBP was targeted to either the plasma membrane (**a**) or Golgi membrane (**b**), and its binding partner FRB-mVenus was fused to Nb6B9. Upon addition of rapalog (rapamycin analog), FKBP and FRB heterodimerize and sequester Nb6B9 to either membranes, disrupting G protein coupling to the receptor and thus blocking signaling from each respective location. **c**. Representative confocal images of HEK293 cells expressing either plasma membrane (PM) or Golgi targeted FKBP showing mVenus-FRB Nb6B9 localization at indicated times after rapalog addition. Representative cells confirm inducible sequestration of Nb6B9 to either PM or Golgi. Arrow indicates PM; Arrowhead indicates Golgi. **d**. Forskolin-normalized D1DR-mediated cAMP response with and without rapalog pretreatment (1μM, 15 min) and SKF81297 at indicated concentrations in HEK293 expressing PM-FKBP (mean ± s.e.m., *n* = 6 biological replicates, P values of 0.0021 and 0.0015 at 10^−7^ and 10^− 6^, respectively) **e**. Forskolin-normalized D1DR-mediated cAMP response with and without rapalog pretreatment (1μM, 15 min) and SKF81297 at indicated concentrations in HEK293 expressing Golgi-FKBP (mean ± s.e.m., *n* = 6 biological replicates, P values of <0.0001 at 10^−7^ and 10^−6^). **f**. Effect of 1μM and 10μM rapalog on forskolin-mediated cAMP response (*n* = 3 biological replicates)

### Local activation of PKA at the Golgi depends on selective activation of Golgi-localized D1DR

A key downstream effector sensed by cAMP is protein kinase A (PKA). PKA is a holoenzyme, consisting of two catalytic and two regulatory subunits (Figure 5a). There are two PKA types (type I and II) that are anchored to distinct subcellular locations through interactions with distinct A kinase anchoring proteins (66). PKA type II has been shown to localize to the perinuclear/Golgi membranes (67). Binding of cAMP to the PKA regulatory subunit induces rapid dissociation and activation of the PKA catalytic subunit (Figure 5a) (68, 69). To test whether cAMP generation by Golgi-localized D1DR/Gs complex results in the activation of PKA at the perinuclear/Golgi, we utilized a previously described HEK293T knock-in cell line expressing a split fluorescent protein, labeling native PKA catalytic subunit gene with GFP (PKAcat-GFP)(35, 70). Stimulation of HEK293T PKAcat-GFP knock in cell lines expressing D1DR with 10nM SKF81297, a concentration that activates both pools of D1DR (Supplementary Figure 3), resulted in rapid dissociation of PKAcat-GFP from the perinuclear/Golgi membranes (Figure 5b top panel). Quantification of these data shows that stimulation with SKF81297 results in sustained activation of PKA at the perinuclear/Golgi regions (Figure 5c, Supplementary Figure 7). We then asked whether PKAcat dissociation is mediated by the activation of D1DR Golgi pool. Given that HEK293T cells do not express OCT2 transporter (Supplementary Figure 5b) and thus DA cannot be transported to the Golgi membranes, we used DA to specifically activate the plasma membrane pool of D1DR. Importantly, stimulation of HEK293T PKAcat-GFP knock-in cells with 10nM DA, a concentration with similar potency as SKF81297 (Supplementary Figure 5a), did not promote PKAcat dissociation (Figure 5b, lower panel, and c). Together, these data indicate that activation of the D1DR at the Golgi, but not the plasma membrane, results in local PKA activation at the perinuclear/Golgi.

**Figure 5.**
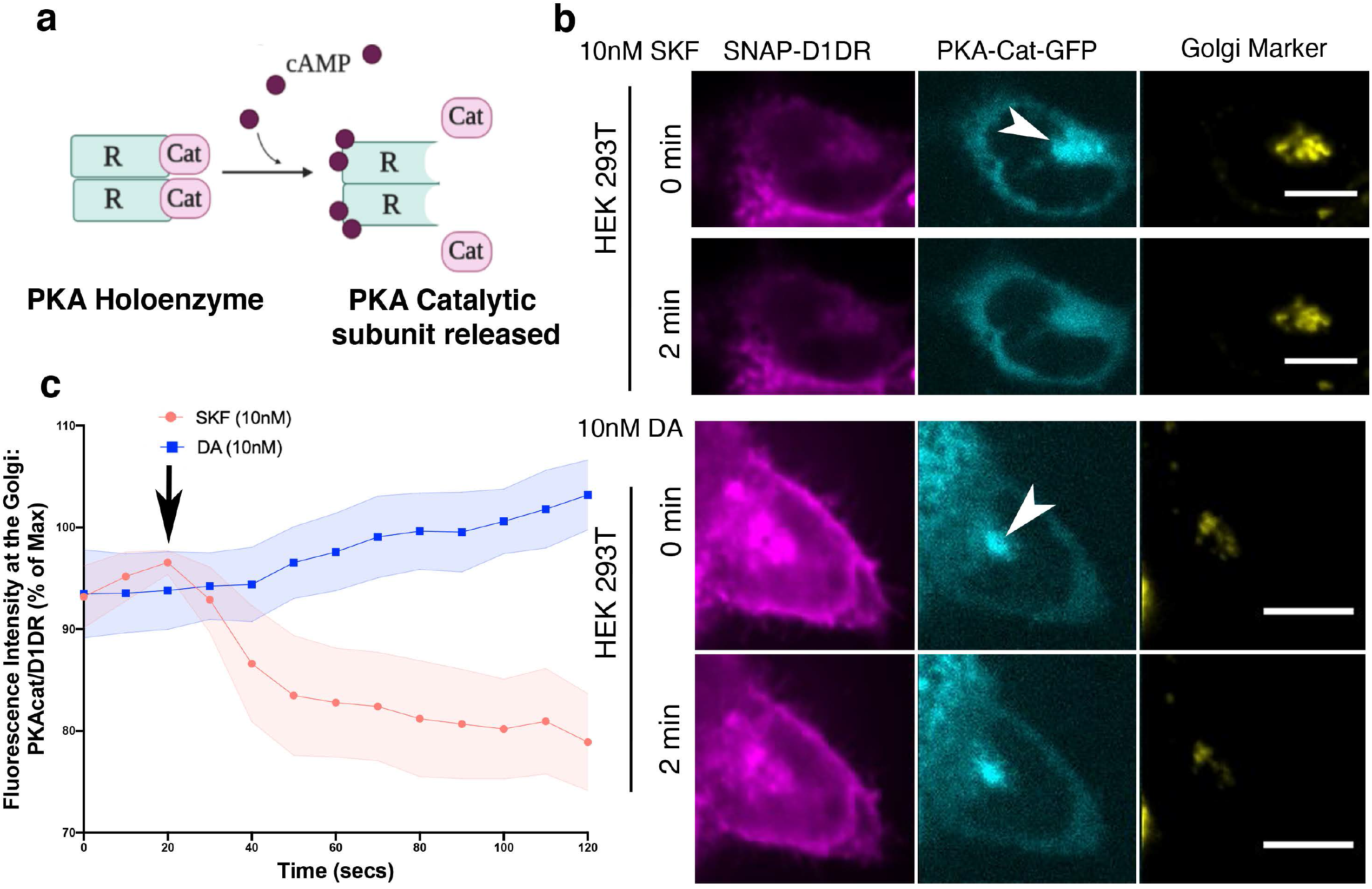
Golgi localized PKA is activated by D1DR at the Golgi. **a**. Model of PKA activation; cAMP binds PKA regulatory subunit (R), rendering PKA catalytic subunit (PKA-cat) dissociation. **b**. Confocal images of representative D1DR-expressing HEK293 cells with endogenous PKA-cat-GFP and GalT-mRFP expression at indicated times after 10nM SKF81927 (top panels; *n* = 11 cells, Pearson’s coefficient 0.53, 3 biological replicates) or 10nM DA (lower panels, *n* = 12 cells, Pearson’s coefficient = 0.64, 3 biological replicates). Arrowhead indicates PKAcat at Golgi membrane; Scale bar = 10μm. **c**. Normalized fluorescence intensity of PKAcat relative to Golgi-D1DR after treatment with 10nM DA or 10nM SKF81927.

### Dopamine uncaging triggers rapid activation of Golgi localized-D1DR and local PKA

To further investigate the role of Golgi-localized D1DR in activating PKA locally, we utilized a photo-sensitive caged dopamine that becomes uncaged upon blue light exposure (Figure 6a). Unlike DA, caged-DA is hydrophobic and thus membrane permeant (59, 71). To ensure that caged-DA accumulates inside the cell and reaches the Golgi-localized D1DR, we incubated HEK293T PKAcat-GFP knock in cells with 1 μM caged-DA for 10 min in a dark incubator. Addition of caged-DA to HEK293T PKAcat-GFP cells did not activate D1DR, as indicated by cytoplasmic localization of Nb6B9-mApple, confirming that DA is inactive in its caged form (Figure 6b top panel). Upon stimulation of cells with blue light for 10 seconds, we observed D1DR activation at the Golgi, as detected by rapid Nb6B9-mApple recruitment to the Golgi membranes within seconds after blue light exposure (Figure 6b and c, Supplementary Movie 9). This was then followed by PKAcat-GFP dissociation from the perinuclear/Golgi regions, as a result of cAMP production and PKA activation (Figure 6b and c, Supplementary Movie 9). These data further support the notion that Golgi-localized D1DR activates PKA locally.

**Figure 6.**
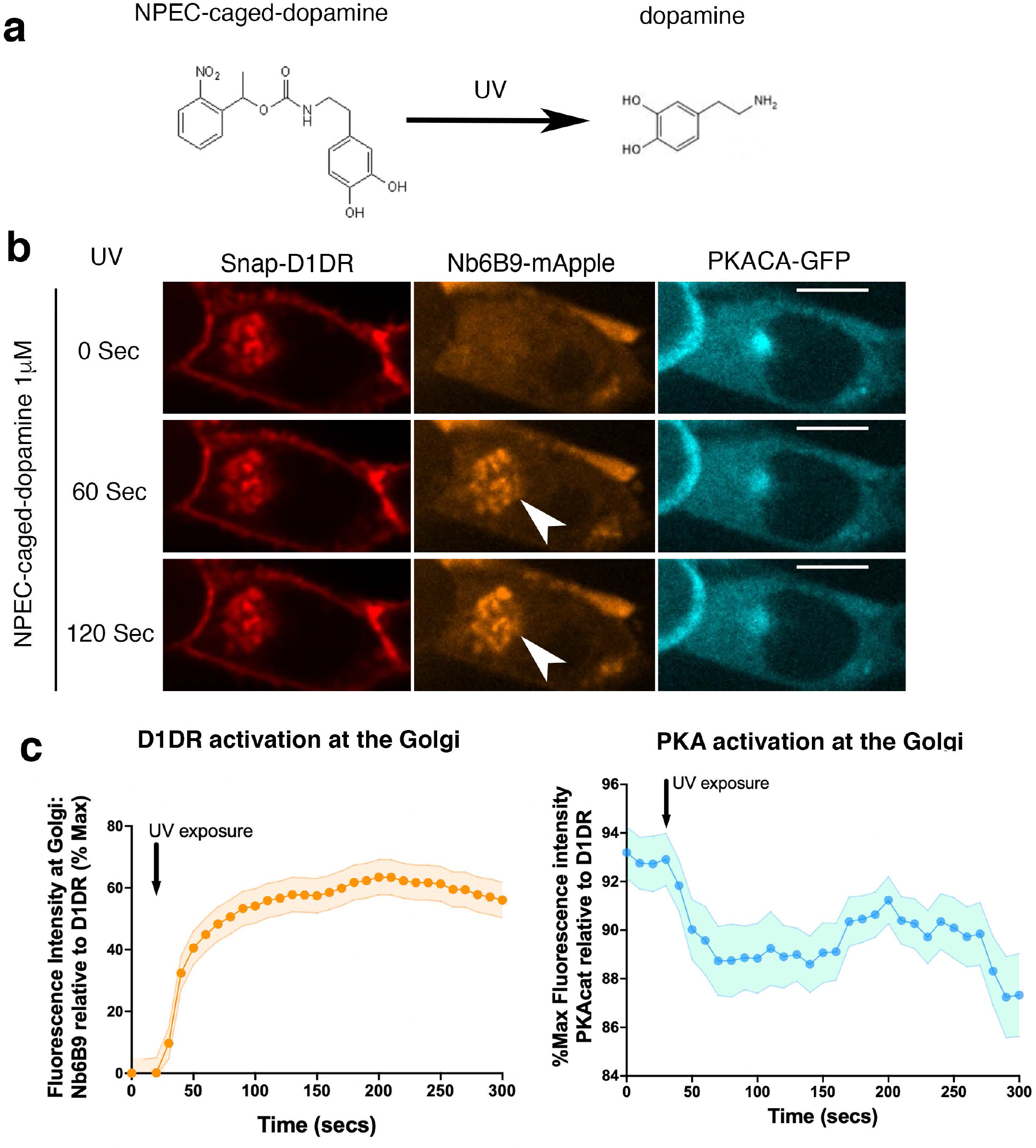
Rapid activation of Golgi localized-D1DR and PKA by photo-release of dopamine. **a**. Dopamine is un-caged from (N)-1-(2 Nitrophenyl) ethylcarboxy-3,4-dihydroxyphenethylamine (NPEC) upon blue light (UV) exposure. **b**. Confocal images of representative D1DR-expressing HEK293 cells with endogenous PKA-cat-GFP and Nb6B9-mApple expression, incubated with 1 μM NPEC-caged dopamine and at indicated times after blue light exposure (*n* = 46 cells, 4 biological replicates). Arrowhead indicates Nb6B9 recruitment to the Golgi membrane; Scale bar = 10μm. **c**. Normalized fluorescence intensity of Nb6B9-mApple and PKAcat relative to Golgi-D1DR after blue light exposure.

## Discussion

Our findings demonstrate for the first time that dopaminergic receptors can signal from the Golgi apparatus. We present evidence that dopamine, a hydrophilic catecholamine, can be transported to the Golgi membrane to reach the pre-existing Golgi pool of D1DRs. This transport is facilitated by OCT2. The Golgi-D1DR comprises a functional signaling pool as it can activate Gαs and stimulate cAMP production. Moreover, we introduced a new approach to selectively interrogate compartmentalized D1DR signaling by inhibiting Gαs coupling, using a nanobody-based chemical recruitment system. Finally, utilizing caged-dopamine, we showed that photo-release of dopamine at the Golgi upon rapid blue light exposure triggers D1DR-mediated cAMP production and local PKA activation.

As the signaling activities of D1DR have thus far been thought to be limited to the plasma membrane, substantial efforts have been focused on designing small molecules agonist of D1DR to *bias* signaling towards a particular signaling pathway, without the consideration of spatial D1DR signaling (16-18, 72). Our findings on D1DR signaling from the Golgi membrane suggest *location-bias* as an overlooked aspect of signaling specificity. The present study demonstrates the important role of local generation of cAMP by GPCRs in controlling local PKA activation at specific subcellular compartments. It is well established that cAMP mediated signaling specificity depends on the function of compartment specific phosphodiesterases, enzymes that degrade cAMP, limiting the diffusion of this second messenger (73-78). Recent measurements of cAMP mobility suggest a nanometer scale diffusion domain (36, 37). The model where cAMP generation by plasma membrane localized receptors propagates in a linear fashion to then control intracellular effectors of cAMP is inconsistent with the nanometer scale of cAMP diffusion range within the cell (37, 38, 79). Thus, our data further provide evidence that PKA activation at a specific compartment requires GPCR activation locally in the vicinity of that compartment. Given that each subcellular membrane compartment has a distinct lipid environment (80), it is likely that PKA activation at each location will recruit a unique set of effectors and proteins and regulates distinct signaling and physiological outcomes. Thus, precisely understanding the functions of the plasma membrane-versus Golgi-localized D1DR in neurons is key for developing better treatments for diseases where dopaminergic signaling is impaired.

Published studies suggest that OCT2 is expressed in a number of tissues and cell types that also express D1DRs (54, 61, 62). Interestingly, however, there are D1DR expressing cell types that do not express OCT2. For instance, we found that within the brain, OCT2 is highly expressed in the striatum and moderately in the cortex. In contrast, OCT2 has little to no expression in the hippocampus or substantia nigra (Supplementary Figure 5c) (54, 56). Consistent with this, we have shown that DA activates Golgi pool of D1DR in MSN cells of the striatum (Figure 3). This could potentially explain some of the distinct cAMP/PKA signaling patterns that have been observed in different D1DR expressing neurons. cAMP-dependent PKA responses have been shown to be long-lasting in striatal neurons compared to pyramidal cortical neurons (59). Therefore, we speculate that the expression pattern of OCT2 and activation of the Golgi-pool of D1DRs may be a determinant of which cell types and tissues exhibit sustained cAMP/PKA signaling.

There are two major DA uptake transport mechanisms: 1) uptake 1 transporters that have high affinity for DA and are mostly localized in presynaptic neurons, and 2) uptake 2 transporters that have low affinity but high capacity for DA and are expressed in various brain regions as well as different organs in the body (50, 57, 81-83). OCT2 belongs to uptake 2 transporter family and has been previously thought to mainly function as an uptake transporter, helping with the clearance of extracellular DA and terminating DA-mediated signaling pathways (53-56). Unlike uptake 1 transporters, uptake 2 transporters can transport catecholamines, including DA, across the membrane, in a bidirectional and electrogenic manner, and independent of Na^+^ and Cl^-^ transport (50, 52). OCTs are localized on both the plasma membrane and subcellular membranes including the outer nuclear membranes near the Golgi (49, 84). Moreover, the resting membrane potential of inner nuclear membrane (∼ -100mV) (85-87) has been reported to be more negative relative to that of the cytoplasmic side of the plasma membrane (∼ -40 to -70 mV). Thus, just as the transport of DA from the extracellular space into the cytoplasm by OCT2 takes advantage of the electrogenic gradient, a similar gradient allows for transport of DA from the cytoplasm across the nuclear envelope which is connected to the lumen of the Golgi membrane.

Accurate measurements of cytoplasmic DA in intact pre-or post-synaptic neurons have been challenging due to lack of sensitivity of most analytical methods and their effects on cell viability (88-90). However, given that DA is present at high millimolar concentrations within the synaptic vesicles(91, 92), it is likely rapid uptake of DA post release will result in high cytoplasmic concentrations. Various Km, ranging from 2 to 46 μM, have been reported for OCT2 transporters (52, 56, 84). Thus, as a low affinity but high-capacity transporter, subcellular OCT2 is likely to encounter high concentrations of cytoplasmic DA under physiological conditions (93). We found that the requirement for OCT2 in activating Golgi-localized D1DRs is seen even at low concentrations of exogenously added DA (10nM) (Supplementary Figure 3). Notably, inhibition of OCT2 but not OCT3 abrogated Golgi-localized D1DR signaling (Figure 2 and 3), highlighting the specificity of OCT2 in this signaling regulation.

The present results expand the concept of GPCR compartmentalized signaling and open additional interesting questions for further studies regarding mechanisms that regulate subcellular activity of other monoamine receptors such as 5-HT (serotonin) and histamine receptors by other monoamine transporters (81, 82). Establishing GPCR signaling from subcellular compartments is the first step in unraveling the physiological consequences of compartmentalized signaling for each GPCR family member.

## Figure legends

**Supplementary Figure 1.**
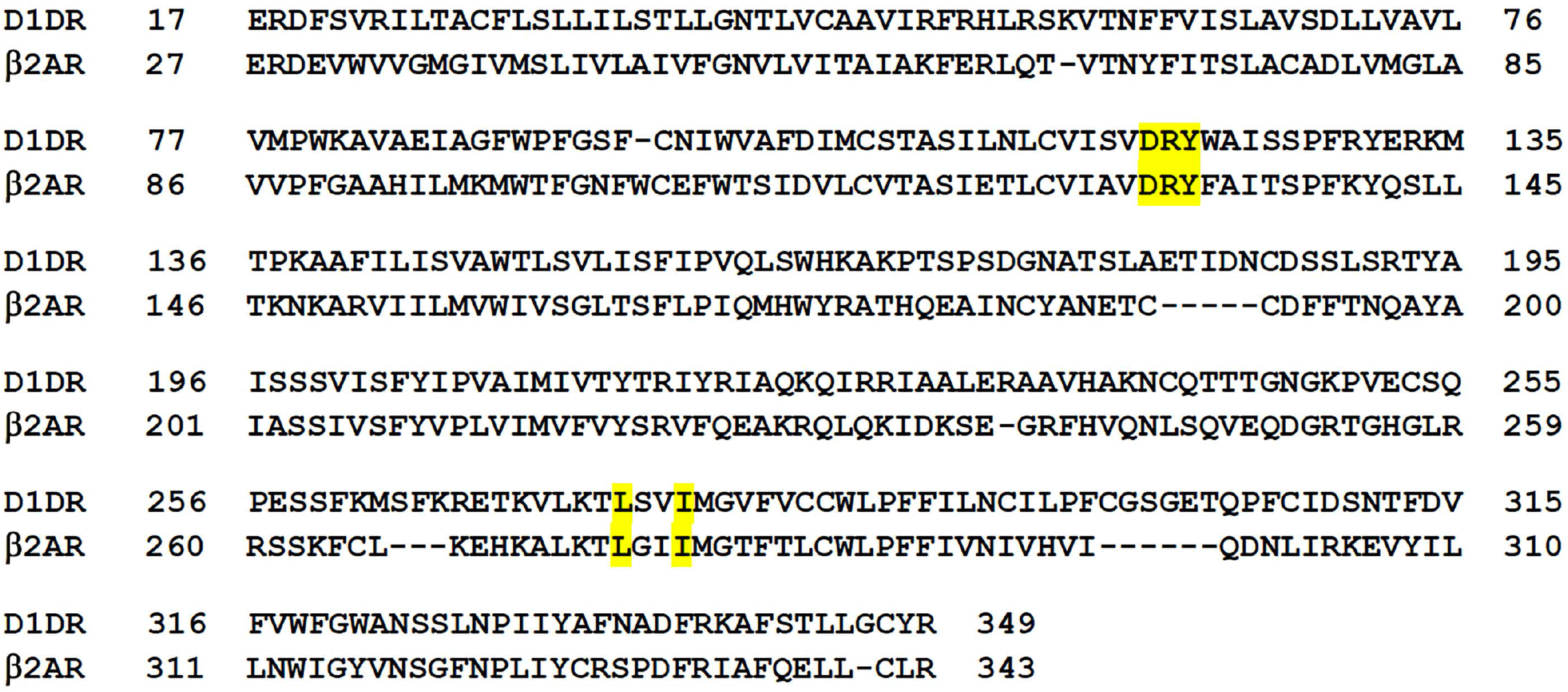
Nanobody binding motifs are conserved between β2AR and D1 dopamine receptor (D1DR).

**Supplementary Figure 2.**
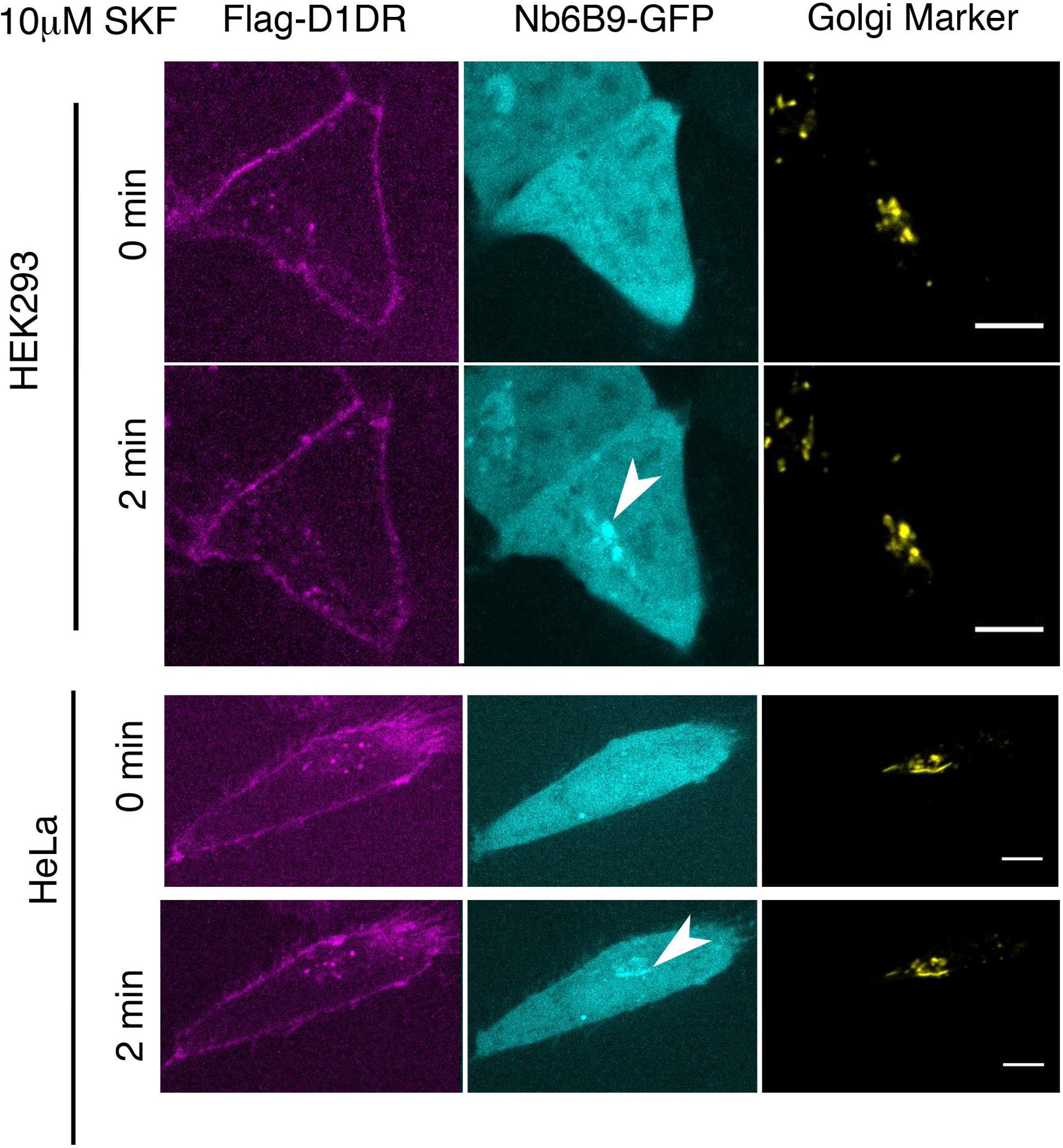
Conformational biosensor detects activated D1DR at the plasma membrane and the Golgi upon SKF81297 stimulation. Representative Flag-D1DR-expressing HEK293 and HeLa cells with Nb6B9-GFP and GalT-mRFP localization at indicated times after 10μM SKF81297 addition. Stimulation with 10μM SKF81297 results in recruitment of Nb6B9 to active D1DR at plasma membrane and Golgi in both HEK 293 and HeLa cells. Scale bar = 10μm.

**Supplementary Figure 3.**
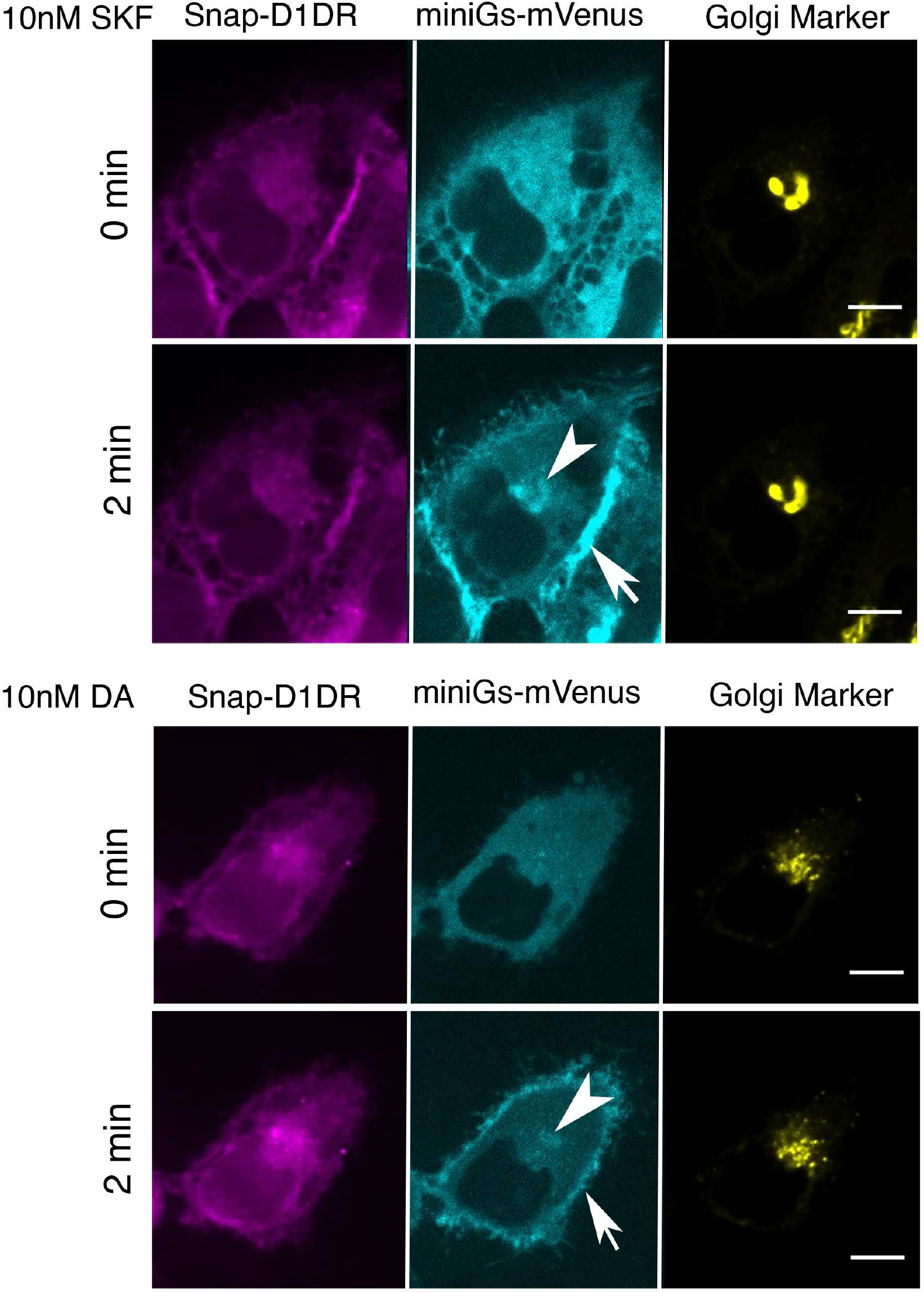
MiniGαs protein biosensor detects active D1DR at the plasma membrane and the Golgi. Representative Snap-D1DR-expressing HeLa cell with miniGs-mVenus and GalT-mRFP localizations at indicated times after 10nM SKF81297 (top panels) and DA (lower panels) addition. Scale bar = 10μm.

**Supplementary Figure 4.**
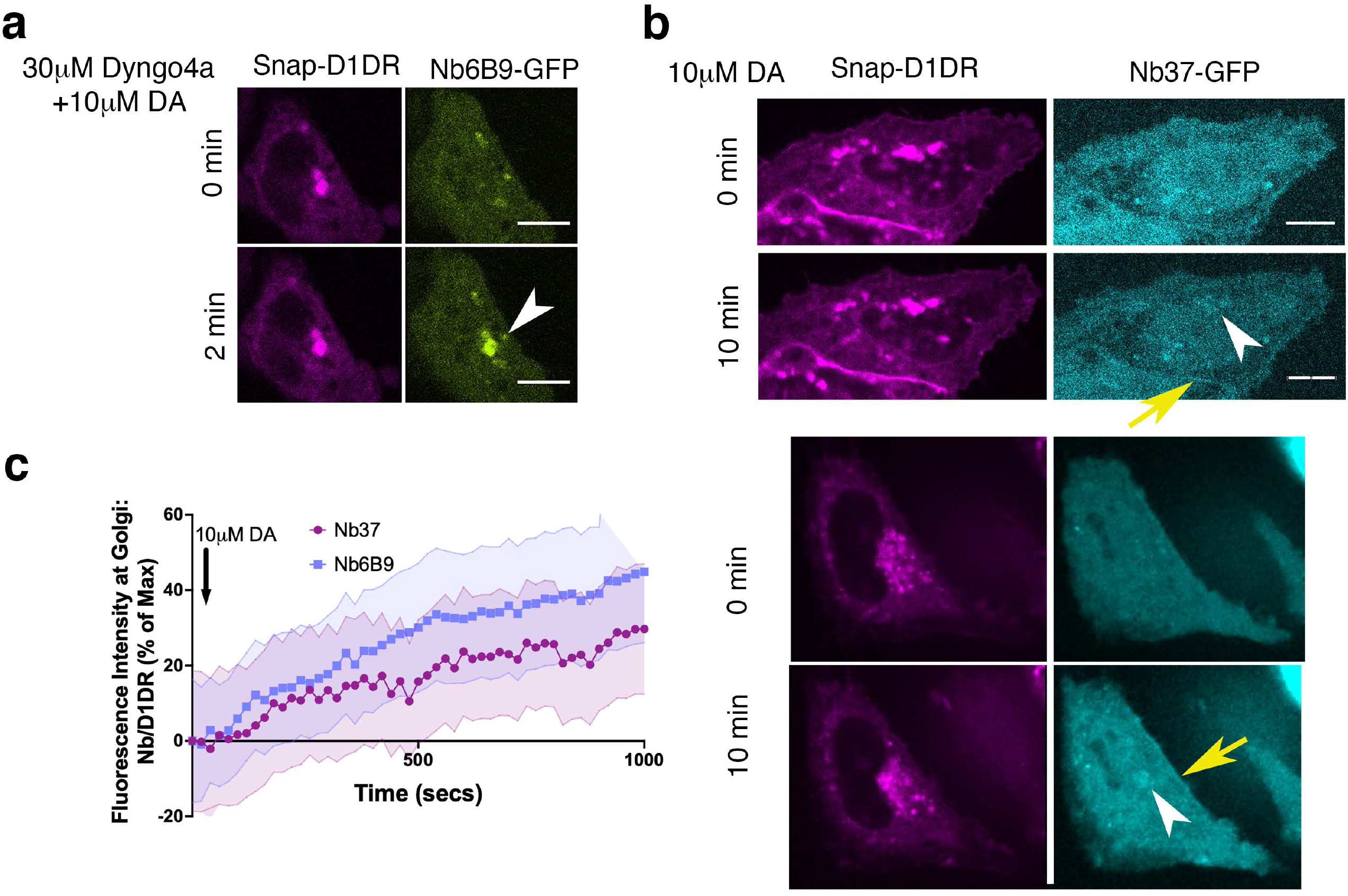
**a**. Representative HeLa cell expressing Snap-D1DR and Nb6B9-GFP. Cells were incubated with 30μM Dyngo-4a, a dynamin inhibitor that blocks endocytosis, stimulated with 10μM DA at indicated times. Golgi-localized D1DR is activated when endocytosis is blocked, suggesting that activation of the Golgi pool is not dependent on D1DR internalization. **b**. Representative HeLa cell expressing Snap-D1DR and Nb37-GFP at indicated times after 10μM DA addition. Nb37 binds to the GPCR-Gs protein complex in the nucleotide free state. Scale bar = 10μm. **c**. Quantification of Nb37-GFP and Nb6B9-GFP intensity at Golgi, normalized to Golgi-D1DR. Both nanobodies show similar kinetics of Golgi-localized D1DR activation after addition of 10μM DA.

**Supplementary Figure 5.**
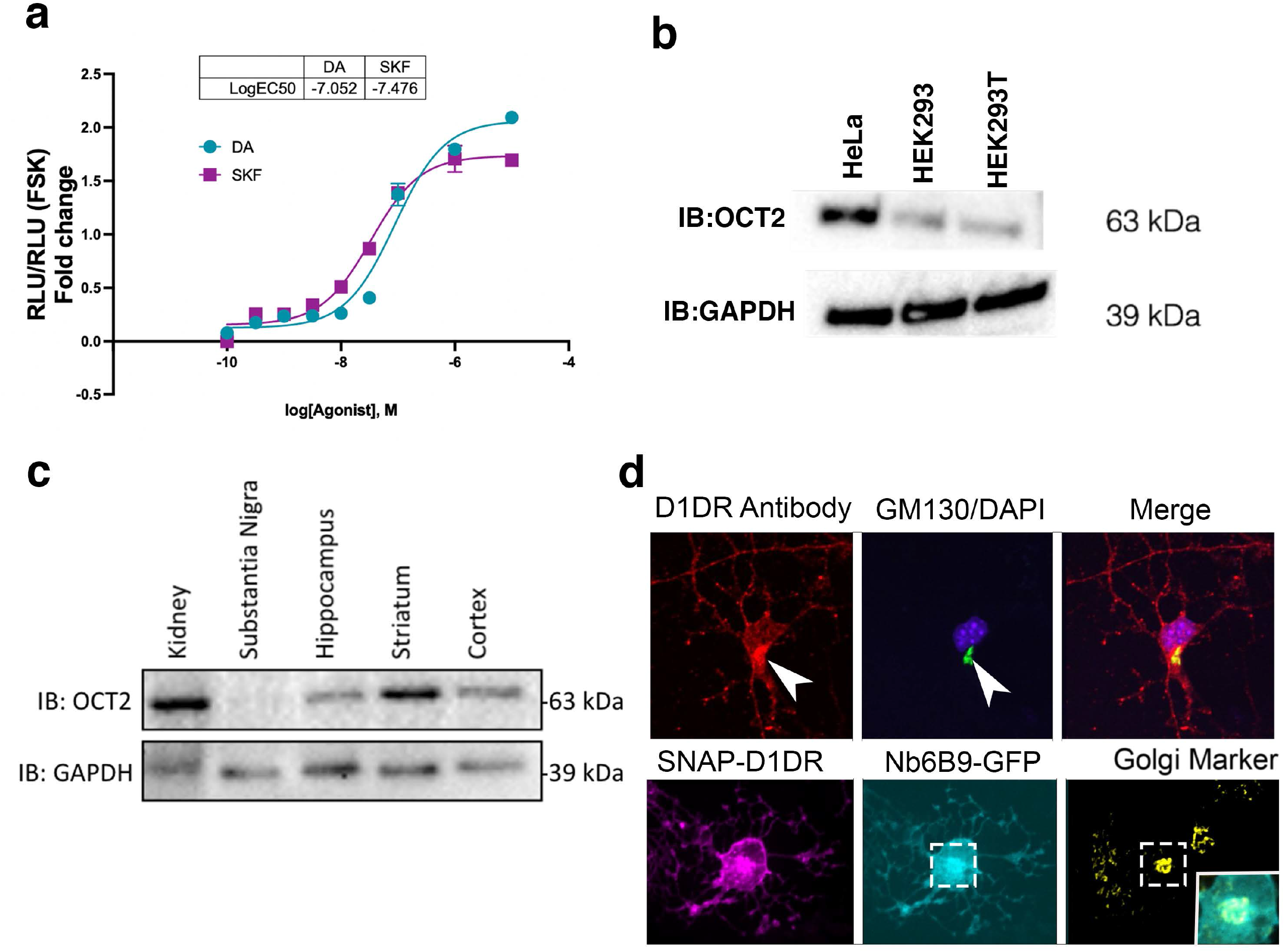
**a**. Dose-response curve of forskolin-normalized D1DR-mediated cAMP response in HEK 293 cells treated with SKF81297 or dopamine; SKF and DA have similar potency and efficacy, thus differences in signaling at Golgi in HEK293 cells are not due to differences in drug potency or efficacy (mean ± s.e.m., *n* = 2 biological replicates). **b**. Detection of OCT2 expression by Western Blot. HeLa cells express OCT2, while HEK 293 cells do not. **c**. Detection of OCT2 expression in different tissue extracts and brain slices by Western Blot. **d**. (Top panel) Endogenous localization of D1DR at the plasma membrane and the Golgi membranes. MSN cells were labeled with D1DR specific antibody and the Golgi antibody (GM130). Arrowhead indicates Golgi localizations. Lower panel, co-localization of Nb6B9-GFP at the Golgi after 10μM DA stimulation for 10 min.

**Supplementary Figure 6.**
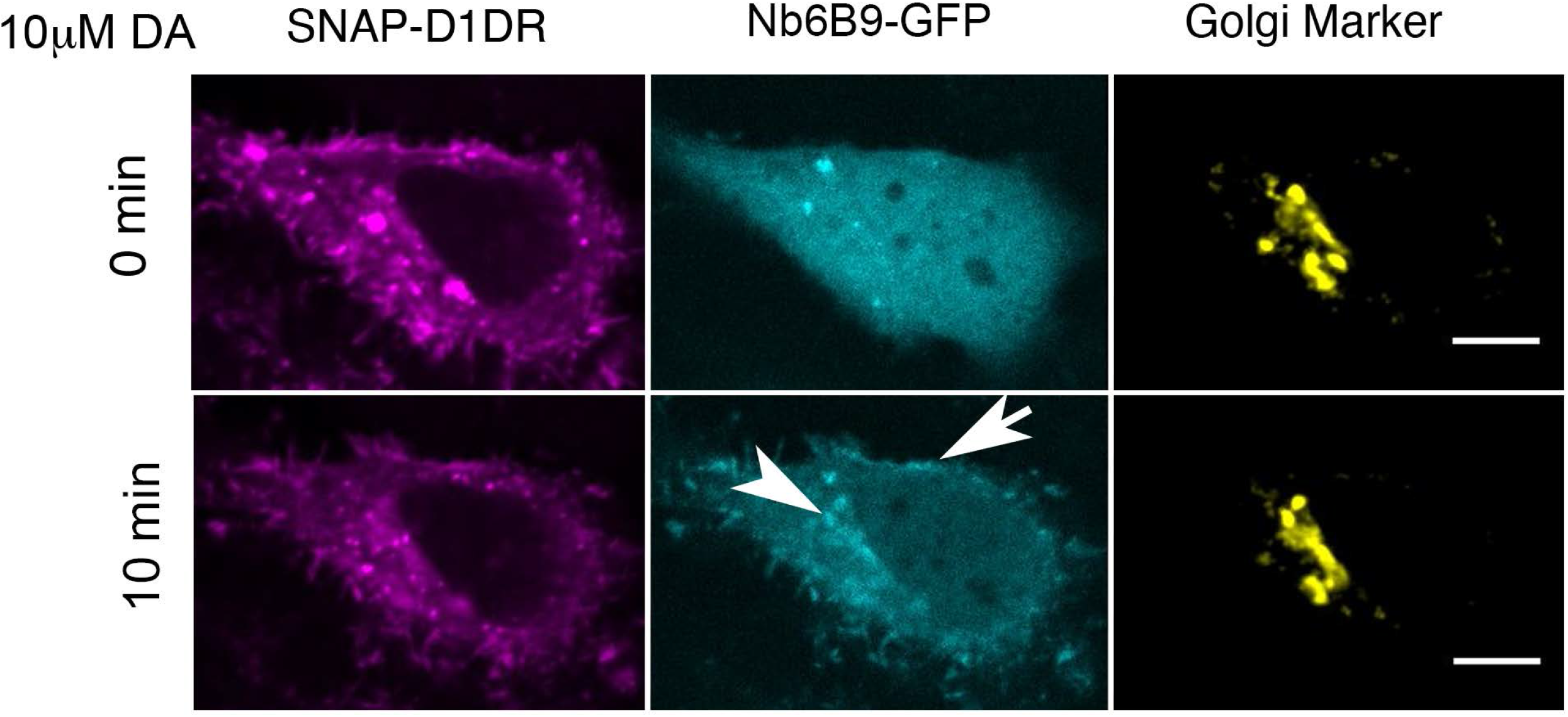
Representative HeLa cell expressing Snap-D1DR, Nb6B9-GFP, and GalT-mRFP pretreated with 10μM corticosterone, an OCT3 selective inhibitor, for 15 min, before and after 10μM dopamine addition. Inhibition of OCT3 does not block Golgi-localized D1DR activation (*n* = 18 cells, Pearson’s coefficient = 0.72, 2 biological replicates).

**Supplementary Figure 7.**
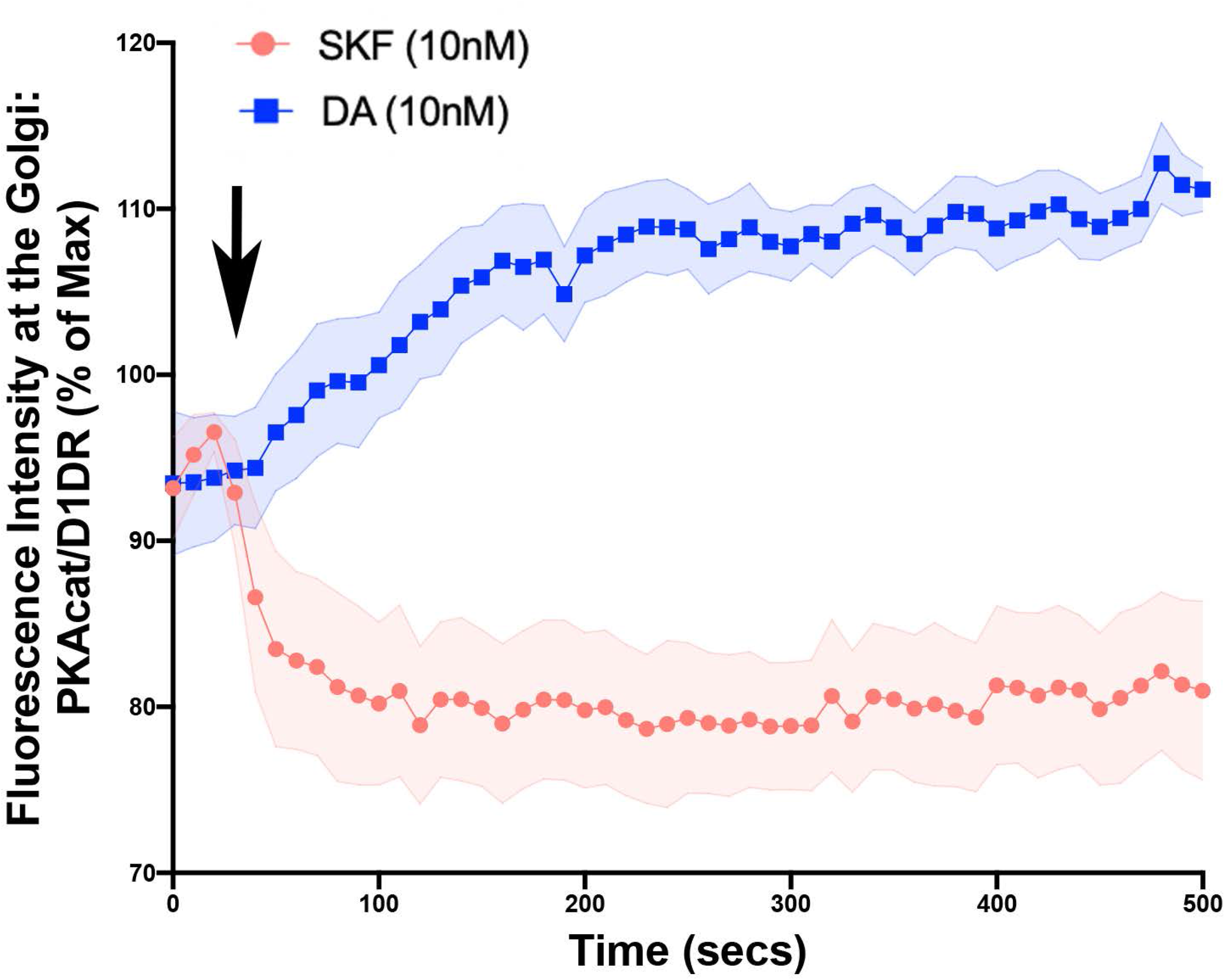
Kinetics of PKAcat-GFP dissociation from the Golgi membrane upon activation of the D1DR Golgi Pools.

**Supplementary Movie 1**. Confocal image series of D1DR expressing HeLa cells (magenta), Nb6B9-GFP (cyan) and the Golgi marker (yellow), incubated with 10μM Dopamine.

**Supplementary Movie 2**. Confocal image series of D1DR expressing HEK293 cells (magenta), Nb6B9-GFP (cyan) and the Golgi marker (yellow), incubated with 10μM Dopamine.

**Supplementary Movie 3**. Confocal image series of D1DR expressing HeLa cells (magenta), Nb6B9-GFP (cyan) and the Golgi marker (yellow), incubated with 10μM SKF81927.

**Supplementary Movie 4**. Confocal image series of D1DR expressing HEK293 cells (magenta), Nb6B9-GFP (cyan) and the Golgi marker (yellow), incubated with 10μM SKF81927.

**Supplementary Movie 5**. Confocal image series of D1DR expressing HEK293 cells (magenta), Nb6B9-GFP (cyan) and OCT2-mApple (yellow), incubated with 10μM Dopamine.

**Supplementary Movie 6**. Confocal image series of D1DR expressing HeLa cells (magenta), Nb6B9-GFP (cyan) and Galt-mApple (yellow), pretreated with 100μM imipramine and incubated with 10μM Dopamine and SKF81297.

**Supplementary Movie 7**. Confocal image series of D1DR expressing MSNs (magenta) and Nb6B9-GFP (cyan), incubated with 10μM Dopamine.

**Supplementary Movie 8**. Confocal image series of D1DR expressing MSNs (magenta) and Nb6B9-GFP (cyan), pretreated with 100μM imipramine and incubated with 10μM Dopamine and SKF81297.

**Supplementary Movie 9**. Confocal image series of D1DR expressing (magenta) HEK293 cells with endogenous PKA-cat-GFP (cyan) and Nb6B9-mApple (orange), incubated with 1μM NPEC-caged-Dopamine.

## Methods

### Cell Culture, cDNA constructs and transfection

HeLa and HEK293 cells (purchased from ATCC as authenticated lines CCL-2, CRL-1573 and CRL 1446 respectively) were grown in Dulbecco’s minimal essential medium supplemented with 10% Fetal Bovine Serum (FBS) without antibiotics. Cell cultures were free of mycoplasma contamination. Signal Sequence-Snap-tagged D1DR was created by amplifying D1DR from Flag-D1DR using 5’-GCCTGGGCTGGGTCTTGGATCCGATGACGCCATGGACG -3’; 5’-ATAGGGCCCTCTAGAGCCTCAGGTTGGGTGCTG -3’ primers, and inserted into the Snap vector using BamHI and XbaI. pVenus-FRB-Nb6B9 was created by amplifying Nb6B9 and FRB from Nb6B9-GFP (40), and pC_4_-R_H_E plasmid (ARIAD Pharmaceuticals), using 5’-TGGTGGACAGGTGCAGCT -3’; 5’-GGATCCTCATGAGGAGACGGTGACCTGGGT -3’ and 5’-

GCTTCGAATTCAATCCTCTGGCAT -3’; 5’-TGCACCTGTCCACCAGCACTA-3 primers respectively, such that it contained the linker sequence GATAGTGCTGGTAGTGCTGGTGGAC, and inserted into the pVenus-C1 vector using EcoRI and BamH1. FKBP-GalT-mApple was created by amplifying FKBP and GalT from KDELr-FKBP and GalT-mCherry plasmids (a generous gift from Dr.Farese lab), using 5’-CATGCTAGCGCCGCCACCATGGGAGTGCAGGTGGAAACCAT-3’, 5’-GAGCTCGAGACCAGCACTACCAGCACTATCCTCCAGCTTCAGCAGCTCCACG3’ and 5’-GCTCAAAGCTTGCCGCCACCGGAAGGCTTCGGGAGCCG-3’, 5’-ACCGGATCCTTAGGCCCCTCCGGTCCGGAGCTCCCCG-3’ primers, respectively and inserted into the pmApple-N1 vector using NheI, XhoI for FKBP and HindIII and BamHI for GalT(40). Transfections were performed using Lipofectamine 2000 (Invitrogen) according to the manufacturer’s instructions. Snap tagged human D1DR constructs were labelled with Snap-cell 647 SiR (New England Biolabs, S9102S) as described previously (94).

### Isolation of murine striatal and hippocampal neurons

Primary striatal neurons were prepared from P1-P2 CD1 pups. In brief, striatum tissues isolated from brains in cold HBSS (w/o Mg^2+^, Ca^2+^ and phenol-red) buffer with 10mM HEPES were treated by HBSS with 0.25%Trypsin and 10mM HEPES buffer at 37°C for 15 min. The digested striatum tissues were rinsed by neural plating media (DMEM with 10%FBS, 30mM HEPES, and PS) twice, and then dissociated by trituration using fire-polished Pasteur pipet in neural plating media. Suspended cells that pass through a 40μm strainer were collected by centrifuging at 350 x g for 5 min. Cells were plated at 10^3^ cells per mm^2^ on the 100μg/ml poly-D-lysine (Sigma) coated imaging dishes or coverslips in neural plating media. After 16-24hr, the culture media were replaced by neural differentiation media (Neural basal media with 10mM Glutamax, B27, and PS). The 50% media were replaced by fresh neural differentiation media every 3-4 days.

### Live-cell confocal imaging

Live cell imaging was carried out using Nikon spinning disk confocal microscope with a 60 ×, 1.4 numerical aperture, oil objective and a CO_2_ and 37 °C temperature-controlled incubator. A 488, 568 nm and 640 Voltran was used as light sources for imaging GFP, mRFP/mApple, and Snap-647 signals, respectively. Cells expressing both Snap-tagged receptor (2μg) and the indicated nanobody–GFP (200ng) were plated onto glass coverslips. Receptors were surface labelled by addition of Snap-Cell 647 SiR (1:1000, New England Biolabs) to the media for 20 min, as described previously. Live cell images where endocytosis was inhibited were carried out by incubating the cells in 30μM Dyngo 4a (ab120689) at 37 °C for 30 minutes before indicated agonist was added. HEK293 PKA-Cat-GFP knock-in cells were a generous gift from the Huang Lab. Indicated agonists (dopamine hydrochloride-Sigma, SKF81297 hydrobromide-Tocris) were added and cells were imaged every 20 s for 20 min in DMEM without phenol red supplemented with 30 mM HEPES, pH 7.4. NPEC-caged-dopamine (Tocris) was incubated for 10 min before cells were stimulated with 3.2 μW/cm^2^ blue light. Time-lapse images were acquired with a CMOS camera (Photometrics) driven by Nikon Imaging Software (NS elements). All live-cell imaging experiments were performed multiple times and on separate days to collect different biological replicates.

### Fixed-cell confocal imaging

Cells were permeabilized with saponin to reduce the cytoplasmic background, as described previously(95). Briefly, MDCK cells were permeabilized with 0.05% saponin (Sigma) in PEM buffer (80mM K-Pipes, pH 6.8, 5 mM EGTA, 1mM MgCl2) for 5 min on ice. Cells were then fixed with 3% paraformaldehyde in PBS for 10 min and then quenched with 50mM Nh4Cl in PBS for 15 min. Primary antibodies (D1DR antibody (ab216644) 1:100, GM130 (BD biosciences 610822) (1:1000), and SLC22A2/OCT2 antibody (ab170871) were diluted in PBS supplemented with 0.05% saponin. Striatal neurons at DIV5 were fixed by 3.7% formaldehyde in PEM buffer for 15 min and then permeabilized by 0.3% Triton in PBS for 5 min at room temperature. D1DR and GM130 antibodies were diluted in TBS with 5% donkey serum and 0.1% Triton X-100. Confocal images were taken using Nikon spinning disk confocal microscope with a 60x 1.4 numerical aperture, oil objective.

### Image analysis and statistical analysis

Images were saved as 16-bit TIFF files. Quantitative image analysis was carried out on unprocessed images using ImageJ software (http://rsb.info.nih.gov/ij). For measuring kinetics of Nb6B9–GFP recruitment at the Golgi membrane over time in confocal images and kinetics of PKA-Cat GFP dissociation from the Golgi, analyses were performed on unprocessed TIFF images using custom scripts written in MATLAB (96). Co-localization analysis at the Golgi was estimated by calculating the Pearson’s coefficient between the indicated image channels with the Golgi marker channel, using the co-localization plug-in for ImageJ (Coloc2). *P* values are from two-tailed unpaired Student’s *t*-tests calculated using Prism 6.0 software (GraphPad Software).

### Luminescence-based cAMP assay

HEK 293 cells stably expressing D1DR were transfected with a plasmid encoding a cyclic-permuted luciferase reporter construct, (pGloSensor-20F, Promega) and luminescence values were measured, as described previously (23). Briefly, cells were plated in 96-well dishes (∼100,000 cells per well) in 500μl DMEM without phenol red/no serum and equilibrated to 37 °C in the SpectraMax plate reader and luminescence was measured every 1.5 min. Software was used to calculate integrated luminescence intensity and background subtraction. In rapamycin heterodimerization experiments, cells were pre-incubated with 1μM A/C heterodimerizer, a rapamycin analog (Takara 635056) for 15 min. 5μM forskolin was used as a reference value in each multi-well plate and for each experimental condition. The average luminescence value (measured across duplicate wells) was normalized to the maximum luminescence value measured in the presence of 5μM forskolin. For rapamycin treated cells, the average luminescence value was normalized to the maximum luminescence value measured in the presence of 5μM forskolin and 1μM Rapamycin.

### Western blotting

Cells from HEK293, HEK293T and HeLa were lysed in extraction buffer (0.2% Triton X-100, 50mM NaCl, 5mM EDTA, 50mM Tris at pH 7.4 and complete EDTA-free protease inhibitor cocktail (Roche). Kidney and neural tissues from B6 adult mice were disrupted in RIPA buffer (50mM Tris at pH 7.4, 150mM NaCl, 1mM EDTA, 1% Triton X-100, 1% Sodium deoxycholate 0.1% SDS and complete EDTA-free protease inhibitor cocktail). After agitation at 4°C for 30 min, supernatants of samples were collected after centrifuging at 15000 x rpm for 10 min at 4°C. Supernatants were mixed with SDS sample buffer for the protein denaturation. The proteins were resolved by SDS-PAGE and transferred to PVDF membrane and blotted for anti-SLC22A2 (ab170871-1:1000) or GAPDH (1:10,000) antibodies to detect OCT2 and GAPDH expression by horseradish-peroxidase-conjugated rabbit IgG, sheep anti-mouse and rabbit IgG (1:10,000 Amersham Biosciences) and SuperSignal extended duration detection reagent (Pierce).

### Data availability

Our research resources, including methods, cells and protocols are available upon request. All reagents developed, such as FRB and FKBP constructs will be available upon request. Data files are included as a excel document. The corresponding author adheres to the NIH Grants Policy and Sharing of Unique Research Resources.

## Acknowledgments

We thank D. Jullie, K. Silm, A. B. Lobingier, A. Manglik. and D. Larsen for assistance, advice and valuable discussion. These studies were supported by the National Institute on General Medicine (GM133521) to R.I.

## Author Contributions

N.P and G.R.R. designed experimental strategy, carried out all of the experiments and analysis. Q.N.M contributed to the cAMP experimental design and analysis. R.I. designed the experimental strategy, contributed to interpreting the results and writing the paper.

## References

1. Sulzer D. How addictive drugs disrupt presynaptic dopamine neurotransmission. Neuron. 2011;69(4):628–49. Epub 2011/02/23. doi: 10.1016/j.neuron.2011.02.010. PubMed PMID: 21338876; PMCID: PMC3065181.

2. Salery M, Trifilieff P, Caboche J, Vanhoutte P. From Signaling Molecules to Circuits and Behaviors: Cell-Type-Specific Adaptations to Psychostimulant Exposure in the Striatum. Biol Psychiatry. 2020;87(11):944–53. Epub 2020/01/14. doi: 10.1016/j.biopsych.2019.11.001. PubMed PMID: 31928716.

3. Di Chiara G, Imperato A. Drugs abused by humans preferentially increase synaptic dopamine concentrations in the mesolimbic system of freely moving rats. Proc Natl Acad Sci U S A. 1988;85(14):5274–8. Epub 1988/07/01. doi: 10.1073/pnas.85.14.5274. PubMed PMID: 2899326; PMCID: PMC281732.

4. Missale C, Nash SR, Robinson SW, Jaber M, Caron MG. Dopamine receptors: from structure to function. Physiol Rev. 1998;78(1):189–225. Epub 1998/02/11. doi: 10.1152/physrev.1998.78.1.189. PubMed PMID: 9457173.

5. Klein MO, Battagello DS, Cardoso AR, Hauser DN, Bittencourt JC, Correa RG. Dopamine: Functions, Signaling, and Association with Neurological Diseases. Cell Mol Neurobiol. 2019;39(1):31–59. Epub 2018/11/18. doi: 10.1007/s10571-018-0632-3. PubMed PMID: 30446950.

6. Felder RA, Seikaly MG, Cody P, Eisner GM, Jose PA. Attenuated renal response to dopaminergic drugs in spontaneously hypertensive rats. Hypertension. 1990;15(6 Pt 1):560–9. Epub 1990/06/01. doi: 10.1161/01.hyp.15.6.560. PubMed PMID: 1971811.

7. Urs NM, Nicholls PJ, Caron MG. Integrated approaches to understanding antipsychotic drug action at GPCRs. Curr Opin Cell Biol. 2014;27:56–62. Epub 2014/04/01. doi: 10.1016/j.ceb.2013.11.002. PubMed PMID: 24680431; PMCID: PMC5702556.

8. Kebabian JW. Dopamine-sensitive adenylyl cyclase: a receptor mechanism for dopamine. Adv Biochem Psychopharmacol. 1978;19:131–54. Epub 1978/01/01. PubMed PMID: 29445.

9. Spano PF, Govoni S, Trabucchi M. Studies on the pharmacological properties of dopamine receptors in various areas of the central nervous system. Adv Biochem Psychopharmacol. 1978;19:155–65. Epub 1978/01/01. PubMed PMID: 358777.

10. Beaulieu JM, Espinoza S, Gainetdinov RR. Dopamine receptors - IUPHAR Review 13. Br J Pharmacol. 2015;172(1):1–23. Epub 2015/02/12. doi: 10.1111/bph.12906. PubMed PMID: 25671228; PMCID: PMC4280963.

11. Honegger KJ, Capuano P, Winter C, Bacic D, Stange G, Wagner CA, Biber J, Murer H, Hernando N. Regulation of sodium-proton exchanger isoform 3 (NHE3) by PKA and exchange protein directly activated by cAMP (EPAC). Proc Natl Acad Sci U S A. 2006;103(3):803–8. Epub 2006/01/13. doi: 10.1073/pnas.0503562103. PubMed PMID: 16407144; PMCID: PMC1334627.

12. Wiederkehr MR, Di Sole F, Collazo R, Quinones H, Fan L, Murer H, Helmle-Kolb C, Moe OW. Characterization of acute inhibition of Na/H exchanger NHE-3 by dopamine in opossum kidney cells. Kidney Int. 2001;59(1):197–209. Epub 2001/01/03. doi: 10.1046/j.1523-1755.2001.00480.x. PubMed PMID: 11135072.

13. Ariano MA, Sortwell CE, Ray M, Altemus KL, Sibley DR, Levine MS. Agonist-induced morphologic decrease in cellular D1A dopamine receptor staining. Synapse. 1997;27(4):313–21. Epub 1998/02/12. doi: 10.1002/(SICI)1098-2396(199712)27:4<313::AID-SYN5>3.0.CO;2-F. PubMed PMID: 9372554.

14. Bloch B, Bernard V, Dumartin B. ‘In vivo’ intraneuronal trafficking of G protein coupled receptors in the striatum: regulation by dopaminergic and cholinergic environment. Biol Cell. 2003;95(7):477–88. Epub 2003/11/05. doi: 10.1016/s0248-4900(03)00080-7. PubMed PMID: 14597266.

15. Vickery RG, von Zastrow M. Distinct dynamin-dependent and -independent mechanisms target structurally homologous dopamine receptors to different endocytic membranes. J Cell Biol. 1999;144(1):31–43. Epub 1999/01/13. doi: 10.1083/jcb.144.1.31. PubMed PMID: 9885242; PMCID: PMC2148123.

16. Undie AS, Weinstock J, Sarau HM, Friedman E. Evidence for a distinct D1-like dopamine receptor that couples to activation of phosphoinositide metabolism in brain. J Neurochem. 1994;62(5):2045–8. Epub 1994/05/01. doi: 10.1046/j.1471-4159.1994.62052045.x. PubMed PMID: 7908949.

17. Panchalingam S, Undie AS. SKF83959 exhibits biochemical agonism by stimulating [(35)S]GTP gamma S binding and phosphoinositide hydrolysis in rat and monkey brain. Neuropharmacology. 2001;40(6):826–37. Epub 2001/05/23. doi: 10.1016/s0028-3908(01)00011-9. PubMed PMID: 11369036.

18. Jin LQ, Goswami S, Cai G, Zhen X, Friedman E. SKF83959 selectively regulates phosphatidylinositol-linked D1 dopamine receptors in rat brain. J Neurochem. 2003;85(2):378–86. Epub 2003/04/05. doi: 10.1046/j.1471-4159.2003.01698.x. PubMed PMID: 12675914.

19. Calebiro D, Nikolaev VO, Lohse MJ. Imaging of persistent cAMP signaling by internalized G protein-coupled receptors. J Mol Endocrinol. 2010;45(1):1–8. Epub 2010/04/10. doi: 10.1677/JME-10-0014. PubMed PMID: 20378719.

20. Ferrandon S, Feinstein TN, Castro M, Wang B, Bouley R, Potts JT, Gardella TJ, Vilardaga JP. Sustained cyclic AMP production by parathyroid hormone receptor endocytosis. Nat Chem Biol. 2009;5(10):734–42. Epub 2009/08/25. doi: 10.1038/nchembio.206. PubMed PMID: 19701185; PMCID: PMC3032084.

21. Feinstein TN, Yui N, Webber MJ, Wehbi VL, Stevenson HP, King JD, Jr., Hallows KR, Brown D, Bouley R, Vilardaga JP. Noncanonical control of vasopressin receptor type 2 signaling by retromer and arrestin. J Biol Chem. 2013;288(39):27849–60. Epub 2013/08/13. doi: 10.1074/jbc.M112.445098. PubMed PMID: 23935101; PMCID: PMC3784700.

22. Kotowski SJ, Hopf FW, Seif T, Bonci A, von Zastrow M. Endocytosis promotes rapid dopaminergic signaling. Neuron. 2011;71(2):278–90. Epub 2011/07/28. doi: 10.1016/j.neuron.2011.05.036. PubMed PMID: 21791287; PMCID: PMC3417347.

23. Irannejad R, Tomshine JC, Tomshine JR, Chevalier M, Mahoney JP, Steyaert J, Rasmussen SG, Sunahara RK, El-Samad H, Huang B, von Zastrow M. Conformational biosensors reveal GPCR signalling from endosomes. Nature. 2013;495(7442):534–8. Epub 2013/03/22. doi: 10.1038/nature12000. PubMed PMID: 23515162; PMCID: PMC3835555.

24. Irannejad R, von Zastrow M. GPCR signaling along the endocytic pathway. Curr Opin Cell Biol. 2014;27:109–16. Epub 2014/04/01. doi: 10.1016/j.ceb.2013.10.003. PubMed PMID: 24680436; PMCID: PMC4968408.

25. Irannejad R, Tsvetanova NG, Lobingier BT, von Zastrow M. Effects of endocytosis on receptor-mediated signaling. Curr Opin Cell Biol. 2015;35:137–43. Epub 2015/06/10. doi: 10.1016/j.ceb.2015.05.005. PubMed PMID: 26057614; PMCID: PMC4529812.

26. Thomsen ARB, Jensen DD, Hicks GA, Bunnett NW. Therapeutic Targeting of Endosomal G-Protein-Coupled Receptors. Trends Pharmacol Sci. 2018;39(10):879–91. Epub 2018/09/06. doi: 10.1016/j.tips.2018.08.003. PubMed PMID: 30180973; PMCID: PMC6508874.

27. Calebiro D, Koszegi Z. The subcellular dynamics of GPCR signaling. Mol Cell Endocrinol. 2019;483:24–30. Epub 2019/01/06. doi: 10.1016/j.mce.2018.12.020. PubMed PMID: 30610913.

28. Lobingier BT, von Zastrow M. When trafficking and signaling mix: How subcellular location shapes G protein-coupled receptor activation of heterotrimeric G proteins. Traffic. 2019;20(2):130–6. Epub 2018/12/24. doi: 10.1111/tra.12634. PubMed PMID: 30578610; PMCID: PMC6387827.

29. Stoeber M, Jullie D, Lobingier BT, Laeremans T, Steyaert J, Schiller PW, Manglik A, von Zastrow M. A Genetically Encoded Biosensor Reveals Location Bias of Opioid Drug Action. Neuron. 2018;98(5):963–76 e5. Epub 2018/05/15. doi: 10.1016/j.neuron.2018.04.021. PubMed PMID: 29754753; PMCID: PMC6481295.

30. Tsvetanova NG, von Zastrow M. Spatial encoding of cyclic AMP signaling specificity by GPCR endocytosis. Nat Chem Biol. 2014;10(12):1061–5. Epub 2014/11/05. doi: 10.1038/nchembio.1665. PubMed PMID: 25362359; PMCID: PMC4232470.

31. Bowman SL, Shiwarski DJ, Puthenveedu MA. Distinct G protein-coupled receptor recycling pathways allow spatial control of downstream G protein signaling. J Cell Biol. 2016;214(7):797–806. Epub 2016/09/21. doi: 10.1083/jcb.201512068. PubMed PMID: 27646272; PMCID: PMC5037407.

32. Jensen DD, Lieu T, Halls ML, Veldhuis NA, Imlach WL, Mai QN, Poole DP, Quach T, Aurelio L, Conner J, Herenbrink CK, Barlow N, Simpson JS, Scanlon MJ, Graham B, McCluskey A, Robinson PJ, Escriou V, Nassini R, Materazzi S, Geppetti P, Hicks GA, Christie MJ, Porter CJH, Canals M, Bunnett NW. Neurokinin 1 receptor signaling in endosomes mediates sustained nociception and is a viable therapeutic target for prolonged pain relief. Sci Transl Med. 2017;9(392). Epub 2017/06/02. doi: 10.1126/scitranslmed.aal3447. PubMed PMID: 28566424; PMCID: PMC6034632.

33. Godbole A, Lyga S, Lohse MJ, Calebiro D. Internalized TSH receptors en route to the TGN induce local Gs-protein signaling and gene transcription. Nat Commun. 2017;8(1):443. Epub 2017/09/07. doi: 10.1038/s41467-017-00357-2. PubMed PMID: 28874659; PMCID: PMC5585343.

34. Jean-Alphonse F, Bowersox S, Chen S, Beard G, Puthenveedu MA, Hanyaloglu AC. Spatially restricted G protein-coupled receptor activity via divergent endocytic compartments. J Biol Chem. 2014;289(7):3960–77. Epub 2014/01/01. doi: 10.1074/jbc.M113.526350. PubMed PMID: 24375413; PMCID: PMC3924264.

35. Peng GE, Pessino V, Huang B, von Zastrow M. Spatial decoding of endosomal cAMP signals by a metastable cytoplasmic PKA network. Nat Chem Biol. 2021;17(5):558–66. Epub 2021/03/03. doi: 10.1038/s41589-021-00747-0. PubMed PMID: 33649598; PMCID: PMC8084946.

36. Bock A, Annibale P, Konrad C, Hannawacker A, Anton SE, Maiellaro I, Zabel U, Sivaramakrishnan S, Falcke M, Lohse MJ. Optical Mapping of cAMP Signaling at the Nanometer Scale. Cell. 2020;182(6):1519–30 e17. Epub 2020/08/28. doi: 10.1016/j.cell.2020.07.035. PubMed PMID: 32846156.

37. Saucerman JJ, Greenwald EC, Polanowska-Grabowska R. Mechanisms of cyclic AMP compartmentation revealed by computational models. J Gen Physiol. 2014;143(1):39–48. Epub 2014/01/01. doi: 10.1085/jgp.201311044. PubMed PMID: 24378906; PMCID: PMC3874575.

38. Agarwal SR, Clancy CE, Harvey RD. Mechanisms Restricting Diffusion of Intracellular cAMP. Sci Rep. 2016;6:19577. Epub 2016/01/23. doi: 10.1038/srep19577. PubMed PMID: 26795432; PMCID: PMC4726171.

39. Richards M, Lomas O, Jalink K, Ford KL, Vaughan-Jones RD, Lefkimmiatis K, Swietach P. Intracellular tortuosity underlies slow cAMP diffusion in adult ventricular myocytes. Cardiovasc Res. 2016;110(3):395–407. Epub 2016/04/20. doi: 10.1093/cvr/cvw080. PubMed PMID: 27089919; PMCID: PMC4872880.

40. Irannejad R, Pessino V, Mika D, Huang B, Wedegaertner PB, Conti M, von Zastrow M. Functional selectivity of GPCR-directed drug action through location bias. Nat Chem Biol. 2017;13(7):799–806. Epub 2017/05/30. doi: 10.1038/nchembio.2389. PubMed PMID: 28553949; PMCID: PMC5733145.

41. Nash CA, Wei W, Irannejad R, Smrcka AV. Golgi localized beta1-adrenergic receptors stimulate Golgi PI4P hydrolysis by PLCepsilon to regulate cardiac hypertrophy. Elife. 2019;8. Epub 2019/08/23. doi: 10.7554/eLife.48167. PubMed PMID: 31433293; PMCID: PMC6726460.

42. Nestler EJ, Luscher C. The Molecular Basis of Drug Addiction: Linking Epigenetic to Synaptic and Circuit Mechanisms. Neuron. 2019;102(1):48–59. Epub 2019/04/05. doi: 10.1016/j.neuron.2019.01.016. PubMed PMID: 30946825; PMCID: PMC6587180.

43. Alburges ME, Hunt ME, McQuade RD, Wamsley JK. D1-receptor antagonists: comparison of [3H]SCH39166 to [3H]SCH23390. J Chem Neuroanat. 1992;5(5):357–66. Epub 1992/09/01. doi: 10.1016/0891-0618(92)90051-q. PubMed PMID: 1358117.

44. Kim YC, Alberico SL, Emmons E, Narayanan NS. New therapeutic strategies targeting D1-type dopamine receptors for neuropsychiatric disease. Front Biol (Beijing). 2015;10(3):230–8. Epub 2015/06/01. doi: 10.1007/s11515-015-1360-4. PubMed PMID: 28280503; PMCID: PMC5340264.

45. Romach MK, Glue P, Kampman K, Kaplan HL, Somer GR, Poole S, Clarke L, Coffin V, Cornish J, O’Brien CP, Sellers EM. Attenuation of the euphoric effects of cocaine by the dopamine D1/D5 antagonist ecopipam (SCH 39166). Arch Gen Psychiatry. 1999;56(12):1101–6. Epub 1999/12/11. doi: 10.1001/archpsyc.56.12.1101. PubMed PMID: 10591286.

46. Rasmussen SG, Choi HJ, Fung JJ, Pardon E, Casarosa P, Chae PS, Devree BT, Rosenbaum DM, Thian FS, Kobilka TS, Schnapp A, Konetzki I, Sunahara RK, Gellman SH, Pautsch A, Steyaert J, Weis WI, Kobilka BK. Structure of a nanobody-stabilized active state of the beta(2) adrenoceptor. Nature. 2011;469(7329):175–80. Epub 2011/01/14. doi: 10.1038/nature09648. PubMed PMID: 21228869; PMCID: PMC3058308.

47. Ring AM, Manglik A, Kruse AC, Enos MD, Weis WI, Garcia KC, Kobilka BK. Adrenaline-activated structure of beta2-adrenoceptor stabilized by an engineered nanobody. Nature. 2013;502(7472):575–9. Epub 2013/09/24. doi: 10.1038/nature12572. PubMed PMID: 24056936; PMCID: PMC3822040.

48. Wan Q, Okashah N, Inoue A, Nehme R, Carpenter B, Tate CG, Lambert NA. Mini G protein probes for active G protein-coupled receptors (GPCRs) in live cells. J Biol Chem. 2018;293(19):7466–73. Epub 2018/03/11. doi: 10.1074/jbc.RA118.001975. PubMed PMID: 29523687; PMCID: PMC5949987.

49. Gasser PJ, Hurley MM, Chan J, Pickel VM. Organic cation transporter 3 (OCT3) is localized to intracellular and surface membranes in select glial and neuronal cells within the basolateral amygdaloid complex of both rats and mice. Brain Struct Funct. 2017;222(4):1913–28. Epub 2016/09/24. doi: 10.1007/s00429-016-1315-9. PubMed PMID: 27659446; PMCID: PMC5362368.

50. Nies AT, Koepsell H, Damme K, Schwab M. Organic cation transporters (OCTs, MATEs), in vitro and in vivo evidence for the importance in drug therapy. Handb Exp Pharmacol. 2011(201):105–67. Epub 2010/11/26. doi: 10.1007/978-3-642-14541-4_3. PubMed PMID: 21103969.

51. Roth M, Obaidat A, Hagenbuch B. OATPs, OATs and OCTs: the organic anion and cation transporters of the SLCO and SLC22A gene superfamilies. Br J Pharmacol. 2012;165(5):1260–87. Epub 2011/10/22. doi: 10.1111/j.1476-5381.2011.01724.x. PubMed PMID: 22013971; PMCID: PMC3372714.

52. Schomig E, Lazar A, Grundemann D. Extraneuronal monoamine transporter and organic cation transporters 1 and 2: a review of transport efficiency. Handb Exp Pharmacol. 2006(175):151–80. Epub 2006/05/26. doi: 10.1007/3-540-29784-7_8. PubMed PMID: 16722235.

53. Taubert D, Grimberg G, Stenzel W, Schomig E. Identification of the endogenous key substrates of the human organic cation transporter OCT2 and their implication in function of dopaminergic neurons. PLoS One. 2007;2(4):e385. Epub 2007/04/27. doi: 10.1371/journal.pone.0000385. PubMed PMID: 17460754; PMCID: PMC1851987.

54. Busch AE, Karbach U, Miska D, Gorboulev V, Akhoundova A, Volk C, Arndt P, Ulzheimer JC, Sonders MS, Baumann C, Waldegger S, Lang F, Koepsell H. Human neurons express the polyspecific cation transporter hOCT2, which translocates monoamine neurotransmitters, amantadine, and memantine. Mol Pharmacol. 1998;54(2):342–52. Epub 1998/08/04. doi: 10.1124/mol.54.2.342. PubMed PMID: 9687576.

55. Bednarczyk D, Ekins S, Wikel JH, Wright SH. Influence of molecular structure on substrate binding to the human organic cation transporter, hOCT1. Mol Pharmacol. 2003;63(3):489–98. Epub 2003/02/28. doi: 10.1124/mol.63.3.489. PubMed PMID: 12606755.

56. Amphoux A, Vialou V, Drescher E, Bruss M, Mannoury La Cour C, Rochat C, Millan MJ, Giros B, Bonisch H, Gautron S. Differential pharmacological in vitro properties of organic cation transporters and regional distribution in rat brain. Neuropharmacology. 2006;50(8):941–52. Epub 2006/04/04. doi: 10.1016/j.neuropharm.2006.01.005. PubMed PMID: 16581093.

57. Grundemann D, Koster S, Kiefer N, Breidert T, Engelhardt M, Spitzenberger F, Obermuller N, Schomig E. Transport of monoamine transmitters by the organic cation transporter type 2, OCT2. J Biol Chem. 1998;273(47):30915–20. Epub 1998/11/13. doi: 10.1074/jbc.273.47.30915. PubMed PMID: 9812985.

58. Tang TS, Bezprozvanny I. Dopamine receptor-mediated Ca(2+) signaling in striatal medium spiny neurons. J Biol Chem. 2004;279(40):42082–94. Epub 2004/08/05. doi: 10.1074/jbc.M407389200. PubMed PMID: 15292232.

59. Castro LR, Brito M, Guiot E, Polito M, Korn CW, Herve D, Girault JA, Paupardin-Tritsch D, Vincent P. Striatal neurones have a specific ability to respond to phasic dopamine release. J Physiol. 2013;591(13):3197–214. Epub 2013/04/05. doi: 10.1113/jphysiol.2013.252197. PubMed PMID: 23551948; PMCID: PMC3717223.

60. Hallett PJ, Spoelgen R, Hyman BT, Standaert DG, Dunah AW. Dopamine D1 activation potentiates striatal NMDA receptors by tyrosine phosphorylation-dependent subunit trafficking. J Neurosci. 2006;26(17):4690–700. Epub 2006/04/28. doi: 10.1523/JNEUROSCI.0792-06.2006. PubMed PMID: 16641250; PMCID: PMC6674081.

61. Double KL, Crocker AD. Dopamine receptors in the substantia nigra are involved in the regulation of muscle tone. Proc Natl Acad Sci U S A. 1995;92(5):1669–73. Epub 1995/02/28. doi: 10.1073/pnas.92.5.1669. PubMed PMID: 7878037; PMCID: PMC42581.

62. Arnsten AF, Cai JX, Steere JC, Goldman-Rakic PS. Dopamine D2 receptor mechanisms contribute to age-related cognitive decline: the effects of quinpirole on memory and motor performance in monkeys. J Neurosci. 1995;15(5 Pt 1):3429–39. Epub 1995/05/01. PubMed PMID: 7751922; PMCID: PMC6578230.

63. Cancino J, Capalbo A, Di Campli A, Giannotta M, Rizzo R, Jung JE, Di Martino R, Persico M, Heinklein P, Sallese M, Luini A. Control systems of membrane transport at the interface between the endoplasmic reticulum and the Golgi. Dev Cell. 2014;30(3):280–94. Epub 2014/08/15. doi: 10.1016/j.devcel.2014.06.018. PubMed PMID: 25117681.

64. Boivin B, Lavoie C, Vaniotis G, Baragli A, Villeneuve LR, Ethier N, Trieu P, Allen BG, Hebert TE. Functional beta-adrenergic receptor signalling on nuclear membranes in adult rat and mouse ventricular cardiomyocytes. Cardiovasc Res. 2006;71(1):69–78. Epub 2006/04/25. doi: 10.1016/j.cardiores.2006.03.015. PubMed PMID: 16631628.

65. Chung KY, Rasmussen SG, Liu T, Li S, DeVree BT, Chae PS, Calinski D, Kobilka BK, Woods VL, Jr., Sunahara RK. Conformational changes in the G protein Gs induced by the beta2 adrenergic receptor. Nature. 2011;477(7366):611–5. Epub 2011/10/01. doi: 10.1038/nature10488. PubMed PMID: 21956331; PMCID: PMC3448949.

66. Soberg K, Skalhegg BS. The Molecular Basis for Specificity at the Level of the Protein Kinase a Catalytic Subunit. Front Endocrinol (Lausanne). 2018;9:538. Epub 2018/09/28. doi: 10.3389/fendo.2018.00538. PubMed PMID: 30258407; PMCID: PMC6143667.

67. Nigg EA, Schafer G, Hilz H, Eppenberger HM. Cyclic-AMP-dependent protein kinase type II is associated with the Golgi complex and with centrosomes. Cell. 1985;41(3):1039–51. Epub 1985/07/01. doi: 10.1016/s0092-8674(85)80084-2. PubMed PMID: 2988780.

68. Tillo SE, Xiong WH, Takahashi M, Miao S, Andrade AL, Fortin DA, Yang G, Qin M, Smoody BF, Stork PJS, Zhong H. Liberated PKA Catalytic Subunits Associate with the Membrane via Myristoylation to Preferentially Phosphorylate Membrane Substrates. Cell Rep. 2017;19(3):617–29. Epub 2017/04/20. doi: 10.1016/j.celrep.2017.03.070. PubMed PMID: 28423323; PMCID: PMC5481286.

69. Walker-Gray R, Stengel F, Gold MG. Mechanisms for restraining cAMP-dependent protein kinase revealed by subunit quantitation and cross-linking approaches. Proc Natl Acad Sci U S A. 2017;114(39):10414–9. Epub 2017/09/13. doi: 10.1073/pnas.1701782114. PubMed PMID: 28893983; PMCID: PMC5625894.

70. Feng S, Sekine S, Pessino V, Li H, Leonetti MD, Huang B. Improved split fluorescent proteins for endogenous protein labeling. Nat Commun. 2017;8(1):370. Epub 2017/08/31. doi: 10.1038/s41467-017-00494-8. PubMed PMID: 28851864; PMCID: PMC5575300.

71. Yapo C, Nair AG, Clement L, Castro LR, Hellgren Kotaleski J, Vincent P. Detection of phasic dopamine by D1 and D2 striatal medium spiny neurons. J Physiol. 2017;595(24):7451–75. Epub 2017/08/07. doi: 10.1113/JP274475. PubMed PMID: 28782235; PMCID: PMC5730852.

72. Kuroiwa M, Bateup HS, Shuto T, Higashi H, Tanaka M, Nishi A. Regulation of DARPP-32 phosphorylation by three distinct dopamine D1-like receptor signaling pathways in the neostriatum. J Neurochem. 2008;107(4):1014–26. Epub 2008/10/01. doi: 10.1111/j.1471-4159.2008.05702.x. PubMed PMID: 18823371.

73. Buxton IL, Brunton LL. Compartments of cyclic AMP and protein kinase in mammalian cardiomyocytes. J Biol Chem. 1983;258(17):10233–9. Epub 1983/09/10. PubMed PMID: 6309796.

74. Agarwal SR, MacDougall DA, Tyser R, Pugh SD, Calaghan SC, Harvey RD. Effects of cholesterol depletion on compartmentalized cAMP responses in adult cardiac myocytes. J Mol Cell Cardiol. 2011;50(3):500–9. Epub 2010/12/01. doi: 10.1016/j.yjmcc.2010.11.015. PubMed PMID: 21115018; PMCID: PMC3049871.

75. Warrier S, Ramamurthy G, Eckert RL, Nikolaev VO, Lohse MJ, Harvey RD. cAMP microdomains and L-type Ca2+ channel regulation in guinea-pig ventricular myocytes. J Physiol. 2007;580(Pt.3):765–76. Epub 2007/02/10. doi: 10.1113/jphysiol.2006.124891. PubMed PMID: 17289786; PMCID: PMC2075464.

76. Steinberg SF, Brunton LL. Compartmentation of G protein-coupled signaling pathways in cardiac myocytes. Annu Rev Pharmacol Toxicol. 2001;41:751–73. Epub 2001/03/27. doi: 10.1146/annurev.pharmtox.41.1.751. PubMed PMID: 11264475.

77. Musheshe N, Schmidt M, Zaccolo M. cAMP: From Long-Range Second Messenger to Nanodomain Signalling. Trends Pharmacol Sci. 2018;39(2):209–22. Epub 2018/01/01. doi: 10.1016/j.tips.2017.11.006. PubMed PMID: 29289379.

78. Gold MG, Gonen T, Scott JD. Local cAMP signaling in disease at a glance. J Cell Sci. 2013;126(Pt 20):4537–43. Epub 2013/10/15. doi: 10.1242/jcs.133751. PubMed PMID: 24124191; PMCID: PMC3795333.

79. Zaccolo M, Zerio A, Lobo MJ. Subcellular Organization of the cAMP Signaling Pathway. Pharmacol Rev. 2021;73(1):278–309. Epub 2020/12/19. doi: 10.1124/pharmrev.120.000086. PubMed PMID: 33334857; PMCID: PMC7770493 this article.

80. Balla T. Phosphoinositides: tiny lipids with giant impact on cell regulation. Physiol Rev. 2013;93(3):1019–137. Epub 2013/08/01. doi: 10.1152/physrev.00028.2012. PubMed PMID: 23899561; PMCID: PMC3962547.

81. Lin Z, Canales JJ, Bjorgvinsson T, Thomsen M, Qu H, Liu QR, Torres GE, Caine SB. Monoamine transporters: vulnerable and vital doorkeepers. Prog Mol Biol Transl Sci. 2011;98:1–46. Epub 2011/01/05. doi: 10.1016/B978-0-12-385506-0.00001-6. PubMed PMID: 21199769; PMCID: PMC3321928.

82. Torres GE, Gainetdinov RR, Caron MG. Plasma membrane monoamine transporters: structure, regulation and function. Nat Rev Neurosci. 2003;4(1):13–25. Epub 2003/01/04. doi: 10.1038/nrn1008. PubMed PMID: 12511858.

83. Reith ME, Zhen J, Chen N. The importance of company: Na+ and Cl-influence substrate interaction with SLC6 transporters and other proteins. Handb Exp Pharmacol. 2006(175):75–93. Epub 2006/05/26. doi: 10.1007/3-540-29784-7_4. PubMed PMID: 16722231.

84. Gasser PJ. Organic Cation Transporters in Brain Catecholamine Homeostasis. Handb Exp Pharmacol. 2021;266:187–97. Epub 2021/05/15. doi: 10.1007/164_2021_470. PubMed PMID: 33987762.

85. Burdakov D, Petersen OH, Verkhratsky A. Intraluminal calcium as a primary regulator of endoplasmic reticulum function. Cell Calcium. 2005;38(3-4):303–10. Epub 2005/08/04. doi: 10.1016/j.ceca.2005.06.010. PubMed PMID: 16076486.

86. Sanchez C, Berthier C, Allard B, Perrot J, Bouvard C, Tsutsui H, Okamura Y, Jacquemond V. Tracking the sarcoplasmic reticulum membrane voltage in muscle with a FRET biosensor. J Gen Physiol. 2018;150(8):1163–77. Epub 2018/06/15. doi: 10.1085/jgp.201812035. PubMed PMID: 29899059; PMCID: PMC6080890.

87. Matamala E, Castillo C, Vivar JP, Rojas PA, Brauchi SE. Imaging the electrical activity of organelles in living cells. Commun Biol. 2021;4(1):389. Epub 2021/03/25. doi: 10.1038/s42003-021-01916-6. PubMed PMID: 33758369; PMCID: PMC7988155.

88. Olefirowicz TM, Ewing AG. Dopamine concentration in the cytoplasmic compartment of single neurons determined by capillary electrophoresis. J Neurosci Methods. 1990;34(1-3):11–5. Epub 1990/09/01. doi: 10.1016/0165-0270(90)90036-f. PubMed PMID: 2259233.

89. Chang Y, Chen Y, Shao Y, Li B, Wu Y, Zhang W, Zhou Y, Yu Z, Lu L, Wang X, Guo G. Solid-phase microextraction integrated nanobiosensors for the serial detection of cytoplasmic dopamine in a single living cell. Biosens Bioelectron. 2021;175:112915. Epub 2021/01/01. doi: 10.1016/j.bios.2020.112915. PubMed PMID: 33383431.

90. Post MR, Sulzer D. The chemical tools for imaging dopamine release. Cell Chem Biol. 2021;28(6):748–64. Epub 2021/04/25. doi: 10.1016/j.chembiol.2021.04.005. PubMed PMID: 33894160; PMCID: PMC8532025.

91. Omiatek DM, Bressler AJ, Cans AS, Andrews AM, Heien ML, Ewing AG. The real catecholamine content of secretory vesicles in the CNS revealed by electrochemical cytometry. Sci Rep. 2013;3:1447. Epub 2013/03/15. doi: 10.1038/srep01447. PubMed PMID: 23486177; PMCID: PMC3596796.

92. Zhang XW, Hatamie A, Ewing AG. Simultaneous Quantification of Vesicle Size and Catecholamine Content by Resistive Pulses in Nanopores and Vesicle Impact Electrochemical Cytometry. J Am Chem Soc. 2020;142(9):4093–7. Epub 2020/02/19. doi: 10.1021/jacs.9b13221. PubMed PMID: 32069039; PMCID: PMC7108759.

93. Wiencke K, Horstmann A, Mathar D, Villringer A, Neumann J. Dopamine release, diffusion and uptake: A computational model for synaptic and volume transmission. PLoS Comput Biol. 2020;16(11):e1008410. Epub 2020/12/01. doi: 10.1371/journal.pcbi.1008410. PubMed PMID: 33253315; PMCID: PMC7728201.

94. Lukinavicius G, Umezawa K, Olivier N, Honigmann A, Yang G, Plass T, Mueller V, Reymond L, Correa IR, Jr., Luo ZG, Schultz C, Lemke EA, Heppenstall P, Eggeling C, Manley S, Johnsson K. A near-infrared fluorophore for live-cell super-resolution microscopy of cellular proteins. Nat Chem. 2013;5(2):132–9. Epub 2013/01/25. doi: 10.1038/nchem.1546. PubMed PMID: 23344448.

95. Lobert VH, Stenmark H. The ESCRT machinery mediates polarization of fibroblasts through regulation of myosin light chain. J Cell Sci. 2012;125(Pt 1):29–36. Epub 2012/01/24. doi: 10.1242/jcs.088310. PubMed PMID: 22266905.

96. Jullie D, Stoeber M, Sibarita JB, Zieger HL, Bartol TM, Arttamangkul S, Sejnowski TJ, Hosy E, von Zastrow M. A Discrete Presynaptic Vesicle Cycle for Neuromodulator Receptors. Neuron. 2020;105(4):663–77 e8. Epub 2019/12/16. doi: 10.1016/j.neuron.2019.11.016. PubMed PMID: 31837915; PMCID: PMC7035187.

